# Ecosystem size and complexity dictate riverine biodiversity

**DOI:** 10.1101/2021.03.08.434467

**Authors:** Akira Terui, Seoghyun Kim, Christine L. Dolph, Taku Kadoya, Yusuke Miyazaki

## Abstract

Larger ecosystems support more species; this ubiquitous pattern is the foundation of current conservation schemes. However, many ecosystems possess a complex spatial structure that cannot be represented by area, and the role of such complexity in scaling biodiversity is largely unknown. Here, we use theory and extensive fish community data from two distinct geographic regions (Japan and United States) to show that ecosystem size and complexity dictate riverine biodiversity. We found that larger and more branched ‘complex’ river networks harbored greater species richness due to increased space and environmental heterogeneity. The complexity effect was comparable to the size effect, and this pattern has emerged regardless of ecological contexts. The dual control of biodiversity may be a pervasive feature that has far-reaching implications for biodiversity conservation.

**One sentence summary:** This study provides the first evidence that ecosystem size and complexity play comparable roles in regulating biodiversity.

## Maintext

Ecologists have long sought to understand the general drivers of biodiversity. One of the most robust empirical generalizations in ecology is the positive relationship between species richness and area, i.e., the species-area relationship (the SAR) (*1*). In 1921, Arrhenius (*2*) formulated the SAR as a power-law *S* = *cA^z^*, an equation currently known as the Arrhenius species-area relationship (*S* is the number of species observed in a given geographic area *A, c* the constant, and *z* the scaling exponent). Since then, the spatial scaling of species richness has been observed in many taxonomic groups (*3*). The SAR is ubiquitous because multiple mechanisms produce an apparently similar pattern. Larger ecosystems typically support more diverse metacommunities due to increased habitat diversity (*4*), larger metacommunity size (*5*), and/or enhanced colonization dynamics (*6*). Importantly, the SAR provides the foundation for global conservation efforts (*7–9*). For example, conservation ecologists have used SAR estimates to design marine and terrestrial protected areas (*7, 8*), which currently encompass more than 30 million km^2^ globally (*10*).

Many ecosystems, however, possess a complex spatial structure that cannot be represented by area – a dimension referred to as scale-invariant complexity (*11, 12*). Such complexity is evident in branching ecosystems, including rivers, trees, and mountain ranges, to name just a few (*12*). Geomorphic or biological processes generate a pronounced self-similarity in complex branching patterns such that the part and the whole look alike (*12*). Even though the branching structure is independent of spatial scale, it forms a physical template that dictates habitat diversity and dispersal corridors for living organisms (*13*). Limited, but accumulating evidence suggests that classical metapopulation and metacommunity theories cannot predict ecological dynamics driven by branching structure (*14–16*), and this recognition has led to recent developments of spatial theories devoted to complex branching ecosystems (*17*). For example, these studies have highlighted key roles of branching structure in driving local biodiversity patterns, such as increased species richness at merging points of branches (*18*). However, most empirical research has explored the consequences of branching complexity for local community structure (*19*) or has relied solely on theoretical arguments with limited replications of branching architecture (*20*). At present, we lack a comprehensive evaluation of how ecosystem complexity controls biodiversity patterns at the metacommunity level. Filling this knowledge gap may provide common ground for achieving successful conservation in spatially complex ecosystems, where accelerated species loss threatens the delivery of ecosystem services (*21*).

Here, we show that ecosystem size and complexity play comparable roles in controlling biodiversity patterns in rivers – a prime example of complex branching ecosystems. Individual streams and rivers flow through different landscapes with distinct geological and climatic backgrounds, serving as a spatial unit of unique in-stream environments (*16, 22–24*). The recurrent merging of diverse tributaries ultimately forms a fluvial network with fractal branching patterns (*12*). As such, the complexity of branching structure, which we define here as the probability of branching per unit river distance (*24, 25*), may represent a macro-scale control of the ecosystem’s habitat heterogeneity (habitat diversity per unit area) (*13, 23, 24*). Meanwhile, ecosystem size (watershed area) should determine the metacommunity size and total habitat diversity (area x heterogeneity), two factors that regulate biodiversity at the metacommunity level (*4, 5*). We predict that ecosystem size and branching complexity increase watershed-scale species richness (γ diversity) by enhancing local species richness (α diversity) and/or spatial difference of species composition (*β* diversity) under different ecological scenarios. The present study combines theory and empirical analysis of extensive community data from two different regions of the globe to provide crucial insights into how ecological communities are structured in complex branching networks.

### Synthesizing ecosystem size and complexity influences

First, we developed a theoretical framework synthesizing the influences of ecosystem size and complexity on biodiversity patterns in branching ecosystems. We depicted branching ecosystems as a spatial network of connected habitat patches which local communities inhabit (*24, 25*) (**Figure 1A and B**). Two factors determine mean environmental conditions at each habitat patch: (1) headwater environments (the most upstream habitat patch) and (2) local environmental noise. Environmental values at headwaters are drawn randomly from a normal distribution and propagate downstream with local environmental noise (i.e., a spatial autoregressive process with white noise). These environmental values recurrently ‘mix’ at confluences considering the relative size of joining tributaries (see **Materials and Methods**). Therefore, our network-generation procedure resembles natural processes of how branching river networks create diverse habitats in a metacommunity. We then simulated metacommunity dynamics in the theoretical branching networks using a general metacommunity model (*26*). In this model, 50 species with different abiotic niches (optimum and width) disperse along a network and compete for resources with varied strengths in temporally dynamic environments (see **Materials and Methods** for model details).

**Figure 1:**
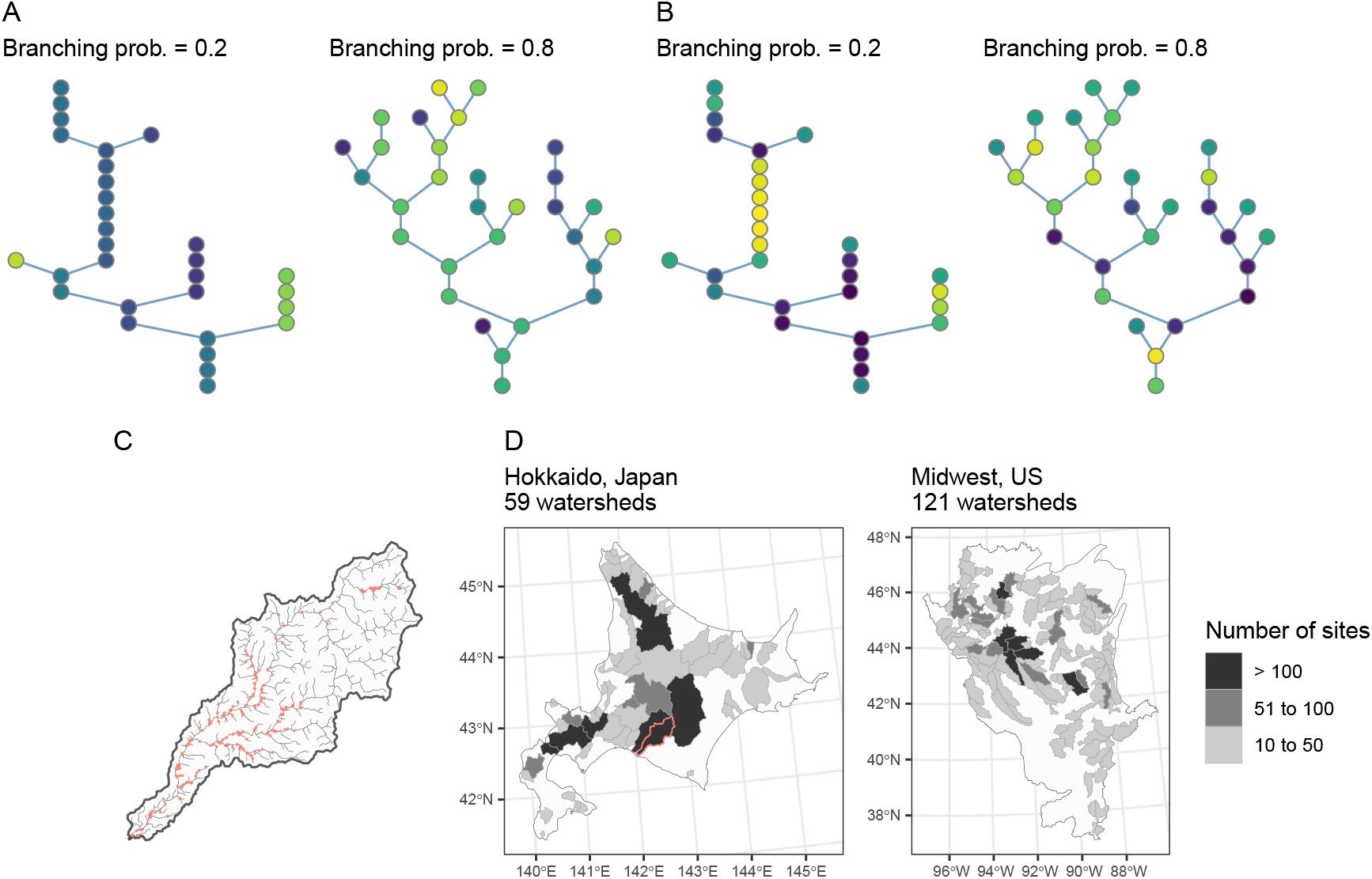
(A and B) Theoretical branching networks generated under contrasting landscape scenarios. Branching river networks are depicted as a network of connected habitat patches, in which the number of habitat patches *N_p_* and branching probability *P_b_* dictate the ecosystem size and complexity (*N_p_* = 30 and *P_b_* = {0.2, 0.8} in this example). Environmental conditions at headwaters (i.e., the most upstream patches) are drawn randomly from a normal distribution and propagate downstream with local environmental noise (Methods). Habitat patches are colored in proportion to environmental values (similar colors have similar environmental values). Panels A and B show distinct landscape scenarios. Environmental variation at headwaters *σ_h_* exceeds the degree of local environmental noise *σ_l_* in A (*σ_h_* = 1, *σ_l_* = 0.01), while the opposite is true in B (*σ_h_* = 0.01, *σ_l_* = 1). (C) Example of intensively surveyed watersheds in Hokkaido, Japan (the red-colored watershed in D). Red dots indicate sampling sites for fish surveys. (D) Map of study regions (left, Hokkaido, Japan; right, Midwest, US). Watersheds (i.e., metacommunities) are gray-shaded in proportion to the number of sampling sites.

Given the results of extensive sensitivity analysis (**Supplementary Text**), we considered 32 simulation scenarios comprising a combination of four landscape and eight ecological scenarios. We distinguished landscape scenarios by setting four combinations of environmental variation at headwaters (*σ_h_* = 0.01, 1) and the degree of local environmental noise (*σ_l_* = 0.01, 1), both of which are defined as standard deviations of normal distributions. When *σ_h_* > *σ_l_*, branching produces greater habitat heterogeneity because headwaters are the primary source of environmental variation (**Figure 1A**). This landscape scenario should reproduce natural patterns of habitat heterogeneity, in which environmental conditions differ greatly among tributaries but are highly correlated within a tributary (*16, 22, 23*). In the meantime, when *σ_h_* ≤ *σ_l_*, local environmental noise masks environmental variation among tributaries, leading to limited influences of branching on habitat heterogeneity (**Figure 1B**). This scenario may reflect human-modified landscapes where the physical or biological distinctiveness of tributaries has been compromised due to human activities (*24, 27*). Thus, the inequality between *σ_h_* and *σ_l_* creates contrasting patterns of habitat heterogeneity within a network. We also considered eight ecological scenarios that differ in dispersal distance (controlled by the rate parameter of an exponential dispersal kernel Θ), dispersal probability *p_d_*, and the maximum value of interspecific competition strength *a_max_* (see **Materials and Methods**). These simulation scenarios were capable of reproducing common spatial patterns of local species richness in rivers, corroborating the appropriateness of our choice in parameter combinations (**Supplementary Text**). Under each simulation scenario, we simulated 1400 time steps of metacommunity dynamics (including 400 time steps of initialization and burn-in periods) in 1000 branching networks with the gradients of ecosystem size (the number of habitat patches: 10 to 150) and complexity (branching probability: 0.01 to 0.99).

Our theoretical analysis yielded results consistent with our prediction. Ecosystem size and complexity both increased γ diversity under a realistic landscape scenario (**Figures 2 and 3**), where the environmental variation at headwaters *σ_h_* is greater than the degree of local environmental noise *σ_l_* (**Figure 1A**). The relationships had a characteristic of power-law (i.e., linear in a log-log scale) and were consistent under various ecological scenarios (compare panels in **Figures 2 and 3**). Importantly, the impact of ecosystem complexity was comparable to that of ecosystem size. Hence, regardless of ecological scenarios, ecosystem size and complexity are likely to be equally important in regulating γ diversity.

**Figure 2:**
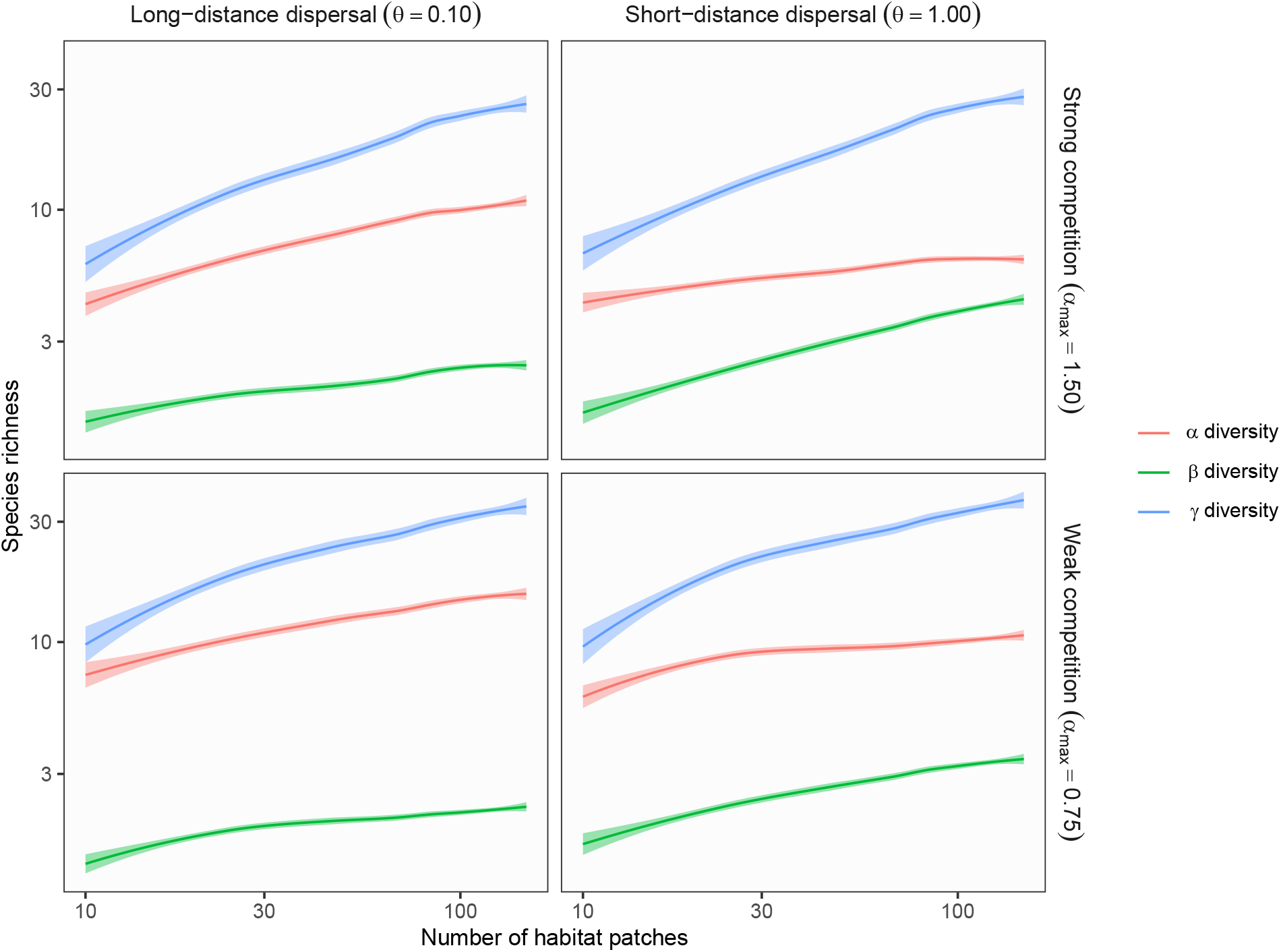
Theoretical predictions for ecosystem size influences (the number of habitat patches) on *α, β,* and *γ* diversity in branching networks. Lines and shades are loess curves fitted to simulated data and its 95% confidence intervals. Each panel represents different ecological scenarios under which metacommunity dynamics were simulated. Rows represent different competition strength. Competitive coefficients (*α_ij_*) were varied randomly from 0 to 1.5 (top, strong competition) or 0.75 (bottom, weak competition). Columns represent different dispersal scenarios. Two dispersal parameters were chosen to simulate scenarios with long-distance (the rate parameter of an exponential dispersal kernel *θ* = 0.10) and short-distance dispersal (*θ* = 1.0). In this simulation, environmental variability among headwaters (i.e., the most upstream patches), which is expressed as the standard deviation of a normal distribution (*σ_h_* = 1.0), was greater than that of local environmental noise occurring at each habitat patch (*σ_l_* = 0.01). Dispersal probability *p_d_* was 0.01 for all the scenarios.

However, dispersal processes affected mechanisms that underlie the positive effects of ecosystem size and complexity on γ diversity (compare left and right columns in **Figures 2 and 3**). We observed a greater contribution of *β* diversity (defined as 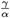) to increased γ diversity when dispersal limitation existed (i.e., species travel short distances). This pattern reflected significant spatial variation in species composition over the branching network and was likely driven by the local association of species’ niche and abiotic environments (i.e., species sorting) (*26, 28*). In contrast, when the dispersal limitation was relaxed (species travel long distances), a clear increase in *α* diversity underpinned the positive relationships between γ diversity and ecosystem properties. The results agree with previous predictions that increased dispersal homogenizes community composition while enhancing local diversity through increased immigrants from suitable habitat patches (i.e., mass effects) (*26, 28, 29*). These patterns were consistent across different levels of dispersal probabilities (**Figures S5 and S12**). The strength of competition decreased the maximum of *α* diversity but did not change the scaling relationships with ecosystem properties (**Figures 2 and 3**). In summary, our theory highlights how apparently similar patterns in γ diversity emerge through different ecological pathways.

**Figure 3:**
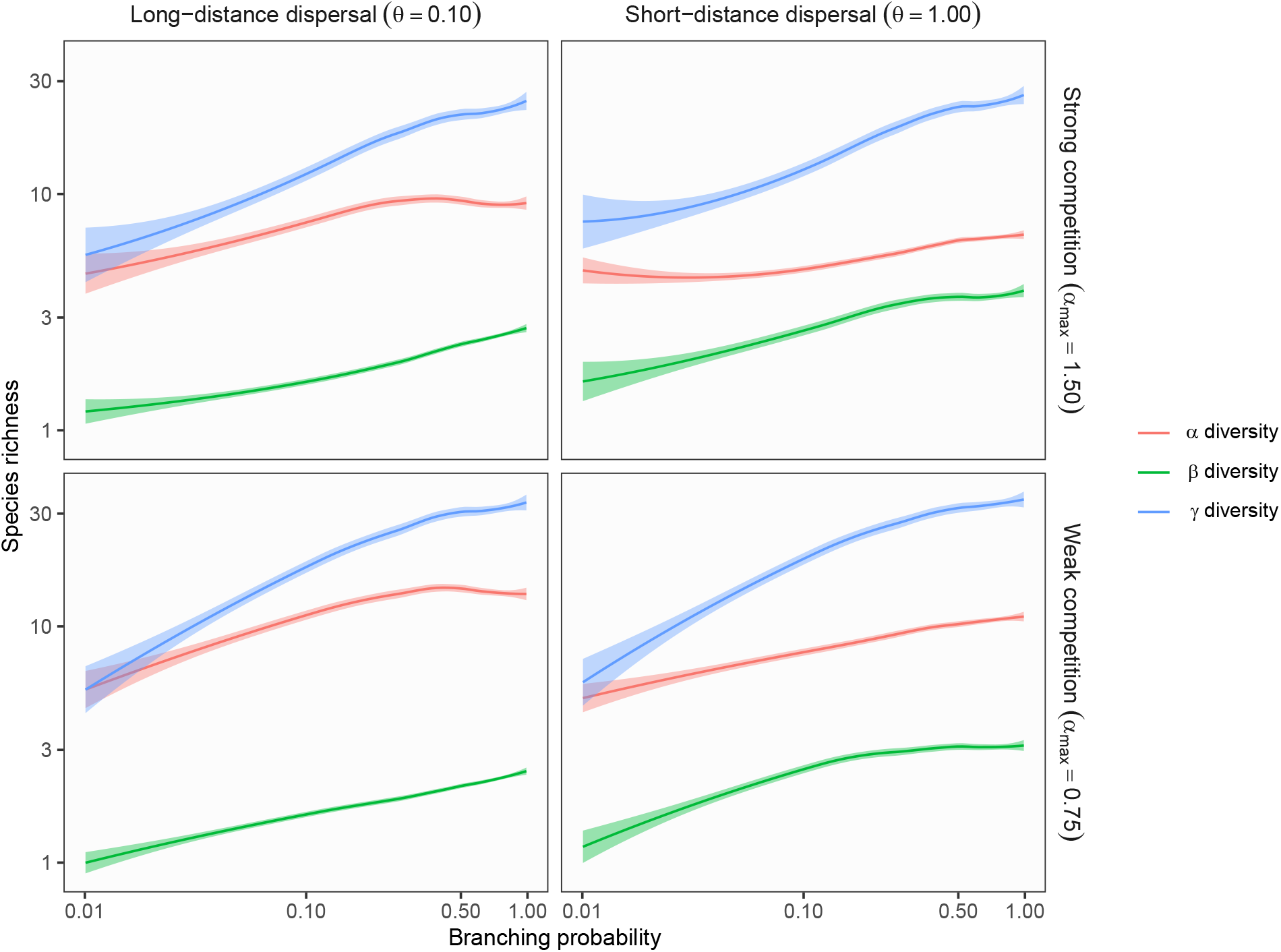
Theoretical predictions for ecosystem complexity influences (branching probability) on *α, β*, and *γ* diversity in branching networks. Lines and shades are loess curves fitted to simulated data and its 95% confidence intervals. Each panel represents different ecological scenarios under which metacommunity dynamics were simulated. See Figure 2 for details.

Influences of ecosystem size and complexity differed significantly in their dependence on landscape scenarios. Ecosystem size had positive effects on γ diversity regardless of landscape scenarios, although the slopes were steeper with greater environmental variation (higher *σ_h_* and/or *σ_l_*) (**Figures 2 and S5-11**). This result is attributable to the fact that larger ecosystems can hold more individuals in a metacommunity (*5*). In contrast, we observed limited or no influences of branching complexity when local environmental noise was equal to or exceeded environmental variation at headwaters (*σ_l_* ≥ *σ_h_*; **Figures S13-18**). Under this scenario, branching has a minor influence on the ecosystem’s habitat heterogeneity because patch-level environmental variation is equivalent to or greater than environmental differences between tributaries (**Figure 1B**). Therefore, this landscape scenario eliminates the positive effect of branching complexity on γ diversity. This theoretical prediction may not apply to pristine or semi-natural river networks where individual streams show distinct and spatially-correlated environmental conditions, including water temperature, water chemistry, and flow/sediment regimes (*22, 23*). Instead, it may be most relevant to severely altered landscapes where localized human disturbance disrupts the environmental distinctiveness of branches through, for example, flow regulations by dams (*27*). Hence, our theory has important implications for riverine biodiversity conservation by pointing to the crucial role of habitat diversity produced by branching structure.

### Empirical evidence from distinct geographic regions

The proposed theory provided important insights into how ecological communities are structured in branching networks; however, empirically testing the predictions is extremely difficult because it requires metacommunity-level replications. To confront this logistical challenge, we compiled fish community data across two geographic regions: Hokkaido Island in Japan and the Midwestern US (Midwest). These regions are located in comparable latitude ranges (**Figure 1D**) but support distinct fish communities (**Tables S5 and S6**). Therefore, this data set provides an excellent opportunity to examine the generality of our theoretical predictions. After careful data selection (see **Materials and Methods** and **Supplementary Text**), we estimated *α, β,* and γ diversity (asymptotic species richness; **Materials and Methods**) at 180 watersheds (59 in Hokkaido and 121 in the Midwest), each of which comprised ≥ 10 sites of presence-absence fish community data (a total of 6605 sites). These watersheds are small enough (40 to 4939 km^2^) to assume that fishes can disperse therein at a multi-generation time scale (*30*), while posing challenges to traverse across watersheds (the ocean or lentic habitats; see **Materials and Methods** for watershed definition). We combined this data set with geospatial information, including watershed characteristics (watershed area and branching probability), climates (annual mean temperature and annual cumulative precipitation), and land use patterns (the fraction of agricultural land use and dam density). Using this data set, we assessed whether observed effects of the macro-scale factors are consistent across the two geographic regions by developing global and region-specific robust regression models. In the global model, we assumed that effects of ecosystem size (watershed area) and complexity (branching probability) are constant across the two regions (i.e., fixed slopes). Meanwhile, the region-specific model assumes region-specific slopes of ecosystem size and complexity (**Materials and Methods**). We compared the support of these competing models using the Bayes factor, a measure of the strength of evidence in favor of one model over the alternative. In our definition (**Materials and Methods**), the Bayes factor of >1 indicates the support for the global model over the region-specific model.

Despite the substantial difference in fish fauna between the study regions, we found strong support for the global model in explaining γ diversity (Bayes factor = 153.8). In line with our theoretical predictions, the estimated γ diversity increased with increasing watershed area (ecosystem size) and branching probability (ecosystem complexity), and these patterns were consistent across the study regions (**Table 1 and Figure 4**). In particular, the effect of ecosystem complexity was striking in its magnitude. Average predictive comparisons (*31*) revealed that an expected increase of γ diversity per 0.1 branching probability was 10.98 ± 4.49 species (**Materials and Methods** and **Supplementary Text**); this level of increase in γ diversity requires 1911 km^2^ in the watershed area (see **Table S7** for the estimated average predictive comparisons). These effects remained significant even after controlling for the potential influences of climates and land use patterns (**Table 1**).

**Figure 4:**
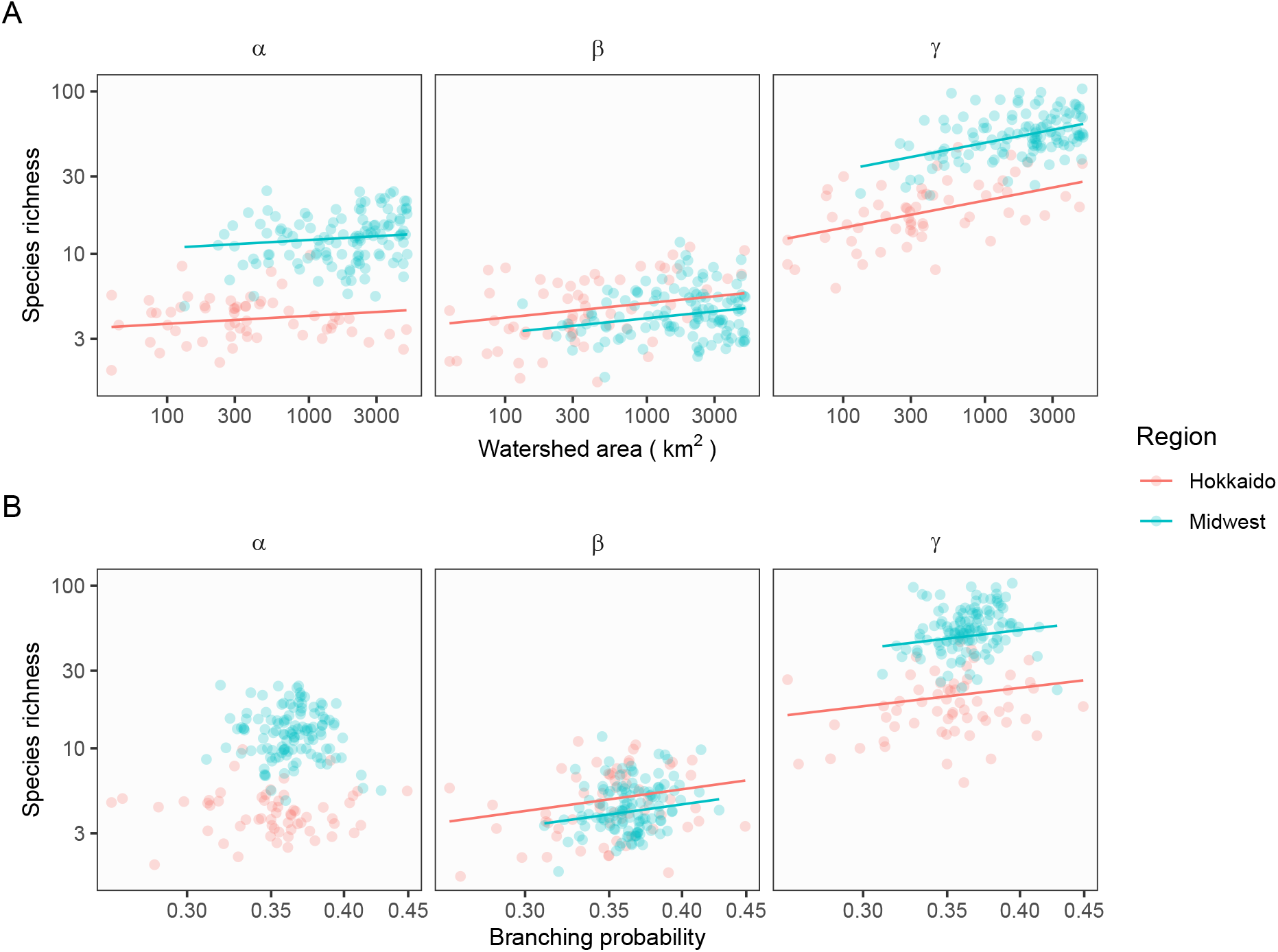
Influences of ecosystem size (A, watershed area) and complexity (B, branching probability) on *a* (left), *β* (middle), and *γ* diversity (right) in Hokkaido and Midwest regions. Dots represent watershed replicates (i.e., metacommunities), and lines are predicted values from the linear regression models.

**Table 1:** Estimated parameters of robust regression models explaining fish species richness in Hokkaido (Japan) and the Midwest (US). The standard errors are shown in parenthesis. Response variables were log-10 transformed. Climate and land use variables (air temperature, precipitation, logit fraction of agriculture, dam density) are deviations from the regional averages and were standardized to a mean of zero and a standard deviation of one prior to the analysis.

|  | Response variable |  |  |
| --- | --- | --- | --- |
| | $\alpha$ diversity | $\beta$ diversity | $\gamma$ diversity |
| $\log_{10}$ Watershed area | 0.05**<br>(0.02) | 0.09***<br>(0.03) | 0.17***<br>(0.02) |
| $\log_{10}$ Branching probability | -0.25<br>(0.30) | 1.07***<br>(0.37) | 0.91***<br>(0.29) |
| Region (Midwest vs. Hokkaido) | 0.47***<br>(0.03) | -0.09***<br>(0.03) | 0.36***<br>(0.02) |
| Air temperature | 0.09***<br>(0.01) | -0.09***<br>(0.02) | 0.01<br>(0.01) |
| Precipitation | -0.03**<br>(0.01) | 0.07***<br>(0.02) | 0.03**<br>(0.01) |
| Logit fraction of agriculture | 0.01<br>(0.01) | 0.01<br>(0.02) | 0.01<br>(0.01) |
| Dam density | 0.004<br>(0.01) | -0.01<br>(0.01) | 0.0003<br>(0.01) |
| Intercept | 0.36**<br>(0.15) | 0.91***<br>(0.19) | 1.23***<br>(0.15) |
| *Note:* *p<0.1; **p<0.05; ***p<0.01 |  |  |  |

Similarly, we found weak to moderate supports for the global models of *α* and *β* diversity (Bayes factor =1.5 and 8.9, respectively). In both regions, *β* diversity responded significantly to ecosystem size and complexity while *α* diversity showing a weaker or a vague response to these ecosystem properties (**Table 1 and Figure 4**). In our simulations, this pattern has emerged under the scenarios with dispersal limitation, which elegantly matches the previous observations of stream fish movement. Field studies (e.g., mark-recapture) recurrently revealed the restricted movement of stream fish, typically limited to several tens to hundreds of meters at an annual scale (*32*). The reciprocal agreement of theoretical and empirical patterns provides indirect but convincing evidence that dispersal limitation, which results in the dominance of species sorting process (*26*), plays a key role in driving the associations between γ diversity and ecosystem properties in rivers.

The consistent effect of branching probability on γ diversity across the study regions is noteworthy because many watersheds in the Midwest have been altered by agricultural land use (mean % agricultural land use: 55% in the Midwest and 6% in Hokkaido). If the intensive land use by humans impairs biological or physical distinctiveness among tributaries, theory predicts a weakened effect of branching probability on γ diversity. However, γ diversity increased significantly with increasing branching probability in this highly modified landscape, suggesting that tributaries still sustain unique environmental conditions to support high spatial variation in species composition. Indeed, *β* diversity increased with increasing branching probability in both regions (**Figure 4**). It is conceivable that local geological and geomorphological differences, such as slope, aspect, and soil porosity, persist in human-dominated landscapes to maintain the diversity of in-stream processes among tributaries. The lack of land use effects (the fraction of agricultural land use) further corroborates our interpretation (**Table 1**). Although our analysis is correlative, the finding is encouraging because the branching complexity of river networks may serve as a natural defense system to human-induced environmental changes.

Overall, our results provide the first empirical evidence that ecosystem size and complexity play comparable roles in controlling biodiversity in rivers. Further, these patterns were fairly consistent with our theoretical predictions that are free from any confounding factors, suggesting that spurious correlations are very unlikely.

### Implications for biodiversity conservation

The emerging complexity-diversity relationship points to several important avenues for riverine biodiversity conservation. First and foremost, there is now a clear need to explicitly consider the dimension of ecosystem complexity to achieve successful conservation. Human-induced habitat alterations, including flow regulation (*27*), habitat fragmentation (*33*), and stream burial (*34*), may compromise or restrict access to the diverse habitats that complex branching networks may support. Hence, it is imperative to recognize the role of branching complexity and minimize the homogenizing effects of human activities. Second, the complexity perspective may provide insights into the spatial planning of conservation efforts. Riverine reserves and local restoration are increasingly recognized as an effective management tool, and designing a spatial network of conservation sites in rivers is an area of active research (*35*). While large reserves or restoration sites are undoubtedly important, our results indicate the potential of coordinated networks of local conservation sites in protecting biodiversity. For example, synergies of multiple small reserves may emerge at the watershed scale when ecologically-distinct tributaries are involved in the design, as evidenced by the recent successful conservation of tropical fishes in Thailand’s Salween basin (*35*).

While the prevailing evidence supports the importance of ecosystem size in scaling biodiversity patterns (*3, 5, 36, 37*), ecosystem complexity has not received the attention it deserves. However, the ubiquity of scale invariant complexity across terrestrial (*11*) and aquatic ecosystems (*12*) calls for more research embracing the two orthogonal dimensions of the ecosystem’s geometric structure. Our fundamental findings of the dual control of biodiversity should apply beyond branching ecosystems because ecosystem size and complexity both represent a physical ‘template’ providing a wide spectrum of niche opportunities for living organisms. Further, the proposed framework may be extended to other aspects of biodiversity; for example, a logical next step is to explore how these ecosystem properties regulate functional diversity (*38, 39*) and shifts in community assembly processes (*40*). Recognizing the dual control of biodiversity broadens viable options for spatial planning of protected areas or restoration sites, thereby helping conserve biodiversity from societal demands that threaten it. The generalization of our findings in spatially complex ecosystems represents a frontier for future research.

## Acknowledgments

We are grateful to numerous people who have been involved in the collection of fish community data. We thank Michio Fukushima and Hirokazu Urabe for providing information on the Hokkaido Freshwater Fish Database HFish and the fish monitoring data at protected watersheds in Hokkaido. We thank Jacques Finlay for his constructive comments on the earlier version of the manuscript. This research was funded by National Science Foundation through a Water Sustainability and Climate Program (WSC) Observatory grant (EAR-1209402): REACH (Resilience under Accelerated Change) and through the Division of Environmental Biology (DEB 2015634), Environmental Protection Agency through a Water Quality Benefits grant (EPA-G2015-STAR-A1), and the University of North Carolina at Greensboro through a start-up fund.

## Author contributions

AT conceived of the project. AT performed theoretical and statistical analysis with inputs from TK. AT, SK, CLD, and YM compiled and managed the fish community data set. AT wrote the first draft while SK, CLD, TK, and YM contributed significantly to the final version.

## Data and code availability

Third parties provided data for empirical data analysis; references are provided where appropriate. R functions for the generation of branching networks and the metacommunity simulation are provided as the R package ‘mcbrnet’ (available at https://github.com/aterui/mcbrnet). Codes for simulations, statistical analysis, figures, and tables are available at https://github.com/aterui/public-proj_stream-diversity.

