## Supplementary materials for "Ecosystem size and complexity dictate riverine biodiversity"

**References** **43**

### Materials and Methods

#### Theory

##### Network generation

We depicted branching ecosystems as a bifurcating network of connected habitat patches (*1*, *2*) (**Figure 1**). Habitat patches, or local communities, can be either non-branching or branching river sections with a unit length $l$, which defines the scale of local species interactions. Two parameters determine the geometric properties of simulated branching networks: the number of habitat patches $N_p$ (ecosystem size) and branching probability $P_b$ (ecosystem complexity). Each habitat patch is assigned to be a branching patch (including upstream terminals) with probability $P_b$ or a non-branching patch with probability $1-P_b$. In this framework, an individual branch is a consecutive series of non-branching patches terminated at a branching patch; therefore, the number of habitat patches in branch $q$, $n_{p,q}$, is a realization of a random variable drawn from a geometric distribution $n_{p,q} \sim Ge(P_b)$ (*1*) but conditional on $\sum_{q=1}^{N_b} n_{p,q} = N_p$ ($N_b$ is the number of branches in a network). The number of branches $N_b$ is identical to the number of branching habitat patches in a network, or the number of 'success' out of $N_p$ trials. This leads to an assumption that $N_b$ is a realization of a random variable drawn from a binomial distribution as $N_b \sim Binomial(N_p, P_b)$. This framework has two merits. First, it reflects observed patterns of branch length distribution, which is known to follow an exponential distribution (a continuous version of a geometric distribution) (*3*). Second, it preserves the fractal nature of branching patterns (*4*). An alternative approach is the use of optimal channel networks (*5*). However, we employed our framework because branching probability $P_b$ is quantifiable in natural river networks, enabling us to compare theoretical predictions with empirical observations.

In our network generation, we first drew $N_b$ from the binomial distribution but selected an odd number to constitute a bifurcating branching network. We then determined $n_{p,q}$ as random draws from the geometric distribution with the constraint of $\sum_{q=1}^{N_b} n_{p,q} = N_p$. The generated branches were organized randomly into a bifurcating branching network.

Two sources of variation characterize the long-term average of the abiotic environment at each habitat patch: environmental variation at headwaters ($\sigma_h$) and local environmental noise ($\sigma_l$). We drew random values from a normal distribution with a mean of zero and a standard deviation of $\sigma_h$ and assigned them to headwater patches (i.e., the most upstream patches). These environmental values at headwaters propagate downstream through a spatial autoregressive process defined as:

$$\mu_{z,k} = \rho \mu_{z,k+1} + \epsilon_k \tag{1}$$

where $\mu_{z,k}$ is the mean environmental value at longitudinal position $k$ ($k$ is the network distance from the outlet patch; $k = 1$ at the outlet), $\rho$ is the strength of spatial autocorrelation and $\epsilon_k$ is the local environmental noise that follows a normal distribution with a mean of zero and a standard deviation of $\sigma_l$. The parameter $\rho$ can take values of 0-1 with larger values indicating greater spatial autocorrelation. In this study, we set $\rho = 1$ to mimic strong spatial autocorrelation in riverine environments. At confluences, we took a weighted mean of environmental values given the relative size of upstream contributing area of joining tributaries:

$$\begin{aligned}\mu_{z,k} &= \omega(\rho\mu^R_{z,k+1} + \epsilon^R_k) + (1-\omega)(\rho\mu^L_{z,k+1} + \epsilon^L_k)\\ \omega &= \frac{A^R_{k+1}}{A^R_{k+1} + A^L_{k+1}}\end{aligned} \tag{2}$$

where $R$ and $L$ denote "right" and "left" branches, respectively, and $A_k$ is the number of upstream habitat patches at longitudinal position $k$ (akin to the upstream watershed area; $A_k = 1$ at the most upstream patch). The parameter $\omega$ takes values of 0-1 and represents the relative influence of the right tributary given the relative size of joining tributaries. With this expression, larger tributaries have a greater influence on the downstream environment, as observed in natural river networks (*6*).

##### Metacommunity model

We simulated metacommunity dynamics in branching river networks following the metacommunity framework proposed by Thompson et al (*7*). In this framework, abiotic environmental conditions (density-independent), intra- and interspecific competition (density-dependent), and dispersal regulate a species' realized population growth. Below, we describe the details of our metacommunity model.

The realized number of individuals $N_{ix}(t+1)$ (species $i$ at patch $x$ and time $t+1$) is given as:

$$N_{ix}(t+1) \sim Poisson(n_{ix}(t) + I_{ix}(t) - E_{ix}(t)) \tag{3}$$

where $n_{ix}(t)$ is the expected number of individuals given the local community dynamics at time $t$, $I_{ix}(t)$ the expected number of immigrants to patch $x$, and $E_{ix}(t)$ the expected number of emigrants from patch $x$. The realized discrete number of individuals is drawn from a Poisson distribution to account for stochasticity in demographic and dispersal processes (*7*, *8*). Local community dynamics are simulated based on the Beverton-Holt equation (*9*):

$$n_{ix}(t) = \frac{N_{ix}(t) r_{ix}(t)}{1 + \frac{r_{0,i}-1}{K_x}\sum_{j=1}^{S}\alpha_{ij}N_{jx}(t)} \tag{4}$$

where $r_{ix}(t)$ is the reproductive rate of species $i$ given the environmental condition at patch $x$ and time $t$, $r_{0,i}$ the maximum reproductive rate of species $i$, $K_x$ the carrying capacity at patch $x$, $\alpha_{ij}$ the competition coefficient between species $i$ and $j$, and $S$ the number of species in a metacommunity. $K_x$ was expressed as a function of the number of upstream habitat patches $A_x$ ($K_x = 100A_x$) to represent increasing habitat size with increasing upstream watershed area (*10*, *11*). The parameter $\alpha_{ij}$ represents the strength of interspecific competition relative to that of intraspecific competition: interspecific competition is greater than intraspecific competition if $\alpha_{ij} > 1$ (intraspecific competition coefficient was set constant as $\alpha_{ii} = 1$). $\alpha_{ij}$ was drawn randomly from a uniform distribution as $\alpha_{ij} \sim Unif(0, \alpha_{max})$. The density-independent reproductive rate $r_{ix}(t)$ is affected by abiotic environments (non-consumable) and determined by the following Gaussian function:

$$r_{ix}(t) = \delta \; r_{0,i} \; exp[-\frac{\{\mu_i - z_x(t)\}^2}{2\sigma^2_{niche,i}}] \tag{5}$$

where $\mu_i$ is the optimal environmental value for species $i$, $z_x(t)$ the environmental value at patch $x$ and time $t$, and $\sigma_{niche,i}$ the niche width of species $i$. We introduced the cost of a wide niche by multiplying $\delta$ (*12*):

$$\delta = exp(-\frac{\sigma^2_{niche,i}}{2\nu^2}) \quad (6)$$

Smaller values of $\nu$ imply greater costs of having a wider niche (i.e., decreased maximum reproductive rate) and reduce the realized reproductive rate $r_{ix}(t)$ near the niche optimum.

The environmental value at patch $x$ and time $t$, $z_x(t)$, is assumed to follow a multivariate normal distribution as $z_x(t) \sim MVN(\mu_z, \Omega_z)$, where $\mu_z$ is the vector of mean environmental conditions of habitat patches and $\Omega_z$ is the variance-covariance matrix. The diagonal elements of the variance-covariance matrix $\sigma^2_z$ represent the degree of temporal environmental variation. The off-diagonal elements $\Omega_{xy}$ represent the spatial autocorrelation in temporal environmental variation as:

$$\Omega_{xy} = \sigma^2_z \ exp(-\phi d_{xy}) \quad (7)$$

$\Omega_{xy}$ denotes the temporal covariance of environmental conditions between patch $x$ and $y$, which is assumed to decay exponentially with increasing distance between the patches $d_{xy}$ (the shortest network path measured as the number of habitat patches). The parameter $\phi$ determines the degree of the distance decay of environmental correlates.

The expected number of emigrants at time $t$ is the product of dispersal probability $p_d$ and $n_{ix}(t)$: $E_{ix}(t) = p_d n_{ix}(t)$. The immigration probability at patch $x$ for species $i$, $\xi_{ix}(t)$, is calculated using the following equation that accounts for separation distance among habitat patches and dispersal capability of a species:

$$\xi_{ix}(t) = \frac{\sum_{y,y\neq x} E_{iy}(t) \ exp(-\theta d_{xy})}{\sum_x \sum_{y,y\neq x} E_{iy}(t) \ exp(-\theta d_{xy})} \quad (8)$$

The parameter $\theta$ regulates the dispersal distance of a species that follows an exponential distribution ($\theta^{-1}$ corresponds to the expected dispersal distance). The expected number of immigrants is calculated as $I_{ix}(t) = \xi_{ix}(t) \sum_x^{N_p} E_{ix}$.

**Simulation**

We used 32 sets of parameter combinations (i.e., simulation scenarios) based on the results of extensive sensitivity analysis (**Supplementary Text**). The parameter combinations comprises four landscape and eight ecological scenarios. Landscape scenarios distinguish spatial patterns of habitat heterogeneity, while ecological scenarios change ecological traits of simulated species. For landscape scenarios, we used four combinations of environmental variation at headwaters ($\sigma_h = 0.01, 1$) and the degree of local environmental noise ($\sigma_l = 0.01, 1$). The inequality of $\sigma_h$ and $\sigma_l$ controls the relationship between branching complexity and habitat heterogeneity. For ecological scenarios, we varied three ecological parameters that are relevant for dispersal and interspecific competition: dispersal distance ($\theta = 0.1$, 1), dispersal probability ($p_d = 0.01$, 0.1), and the maximum value of competition coefficients ($\alpha_{max} = 0.75$, 1.5). This setup results in eight ecological scenarios with different levels of dispersal and competition.

We allowed interspecific variation in niche optimum and width, which were drawn from uniform distributions as $\mu_i \sim Unif(-1, 1)$ and $\sigma_{niche,i} \sim Unif(0.1, 1)$. We used fixed values for the following parameters: maximum reproductive rate ($r_{0,i} = 4$), niche cost ($\nu = 1$), the degree of temporal environmental variation ($\sigma_z = 0.1$), and the extent of spatial autocorrelation in temporal environmental dynamics ($\phi = 0.05$).

Under each simulation scenario, we simulated 1400 time steps of metacommunity dynamics (including 200 time steps for initialization and 200 time steps for burn-in) in 1000 branching networks with the gradients of ecosystem size (the number of habitat patches: 10 to 150) and complexity (branching probability: 0.01 to 0.99). During the first 200 time steps (an initialization period), we initialized the simulation by seeding each habitat patch with populations of each species drawn from a Poisson distribution with a mean of 0.5. We repeated the seeding every 10 time steps during the initialization to allow species to establish their populations if they can grow from low abundances. After the initialization period, we ran 200 time steps as a burn-in period with no seeding to minimize influences of initial conditions. We saved the last 1000 time steps to estimate the temporal averages of $\alpha$, $\beta$, and $\gamma$ diversity. We removed simulation replicates in which no species established populations over the initial 400 time steps (18 replicates out of 32000 replicates). Ecosystem size and complexity were drawn randomly from uniform distributions as: $N_p \sim Unif(10, 150)$ and $P_b \sim Unif(0.01, 0.99)$ ($N_p$ was rounded to the nearest integer before running simulations). Values of simulation parameters were summarized in **Table S2**. We performed simulations using R version 4.0.2 (*13*).

#### Empirical data

##### Fish community data

We compiled fish data at two geographic regions: the Hokkaido island, Japan, and the Midwest, US. For Hokkaido, we used data from the Hokkaido Freshwater Fish Database HFish (*14*, *15*), monitoring data at protected watersheds (*1*, *16*), and primary data collected by the authors (*10*, *17*), which collectively cover the entire island. For the Midwest, we assembled fish community data collected by the Iowa Department of Natural Resources (*18*), Illinois Environmental Protection Agency and Illinois Department of Natural Resources (*19*), Minnesota Pollution Control Agency (*20*), and Wisconsin Department of Natural Resources (*21*). We detailed procedures for initial data selection in **Supplementary Text**.

We considered watersheds as independent replicates of metacommunities if the river network flows into one of the following: the ocean, a large lake/reservoir ($\geq$ 10 km$^2$ in the areal area) or a large river that may represent lentic habitats ($\geq$ 5000 km$^2$ in the watershed area). Metacommunities with $\geq$ 10 sampling sites were used in our analysis because a small sample size inflates statistical uncertainties in estimating diversity metrics (**Supplementary Text**).

We estimated $\alpha$, $\beta$, and $\gamma$ diversity at each watershed. $\gamma$ diversity is the total species richness at each watershed. Since it is impossible to sample all species present in each watershed, we used asymptotic species richness as a proxy for $\gamma$ diversity. Asymptotic species richness can be interpreted as the estimated true species richness in a given area when estimated using spatial replicates of local community data within a metacommunity (*22*). We estimated asymptotic species richness using sampling sites located in the same watershed (i.e., spatial replicates within a metacommunity) via the R package 'iNEXT' (*23*). We assessed the completeness of the observed species richness based on sample coverage, which is the total proportional abundances of observed species to the whole metacommunity (*22*). We excluded watersheds whose sample coverage was below 0.9 because the estimates of asymptotic species richness may contain non-negligible statistical uncertainty. $\alpha$ diversity was estimated by taking an average of local species richness at each site. $\beta$ diversity was estimated by dividing the estimated $\gamma$ diversity by $\alpha$ diversity (*24*).

After the data selection procedure, we obtained diversity estimates from 180 watersheds (Hokkaido: 59, Midwest: 121), which encompassed a total of 6605 sites (Hokkaido: 2607, Midwest: 3998). Observations ranged from 1990 to 2013 in Hokkaido and from 1994 to 2015 in the Midwest. Fish species observed in the final data set were listed in **Tables S5 and S6**.

##### Environmental data

We collected data for watershed characteristics (watershed area [km$^2$], branching probability, and mean elevation [m]), climatic variables (mean annual air temperature [degree Celsius] and cumulative precipitation [mm]), and land use (the fractions of forest, urban, and agriculture, and dam density per watershed area [km$^{-2}$]). We used Hydrologically Adjusted Elevations of MERIT Hydro (*25*) to delineate watershed polygons and river lines (defined as $\geq$ 1 km$^2$ in the watershed area), which were separated by either the ocean, large lakes/reservoirs ($\geq$ 10 km$^2$ in the areal area), or large rivers ($\geq$ 5000 km$^2$ in the watershed area). The ocean and large lakes/reservoirs were extracted from Global 1-second Water Body Map ver 1.0 (*26*). Watershed area was estimated based on the delineated watershed polygons. We estimated branching probability by fitting exponential distributions to histograms of branch length (unit: kilometer, river segment between successive confluences or a confluence and the outlet/upstream terminal) as $branch\ length \sim Exp(\lambda)$. A derived statistical quantity $1 - e^{-\lambda}$ corresponds to the theoretical branching probability $P_b$. We obtained the mean elevations for each watershed polygon using Hydrologically Adjusted Elevations of MERIT Hydro (*25*). We estimated the spatial averages of mean annual air temperature and cumulative precipitation for each watershed (~1 km resolution), which were extracted from WorldClim 1.4 (*27*). We calculated the fractions of forest, urban, and agriculture for each watershed using land use data in 2015 from Copernicus Global Land Service (100-m resolution) (*28*). We calculated dam density per watershed area ($km^{-2}$) based on the Global Reservoir and Dam Database, Version 1 (GRanDv1) (*29*). We used ArcMap 10.7, SAGA 6.3.0, and the following R packages to perform geospatial analysis: 'RSAGA' (*30*), 'sf' (*31*) 'raster' (*32*) 'stars' (*33*), and 'exactextractr' (*34*).

**Statistical analysis**

We developed robust regression models with an M-estimation method to examine the effects of macro-scale drivers on diversity metrics. The robust regression analysis is insensitive to outliers of a response variable; therefore, this method allows us to account for uncertainties of our diversity estimates (see **Fish community data**). Log-transformed diversity metrics ($\alpha$, $\beta$ and $\gamma$ diversity) at watershed $w$, $log_{10}\ y_w$, were assumed to follow a normal distribution as $log_{10}\ y_w \sim Normal(\mu_w, \sigma^2)$. The parameter $\mu_w$ was related to linear predictors, including log-transformed watershed area (ecosystem size; $log_{10}\ WA$), log-transformed branching probability (ecosystem complexity; $log_{10}\ P_b$), region (coded as Hokkaido = 0, Midwest = 1; *region*), air temperature, precipitation, the fraction of agricultural land use (logit-transformed), and dam density. Although our primary focus of this analysis is to assess the effects of watershed area and branching probability, we included climate and land use variables to control their effects statistically. We did not include the fractions of urban and forest land use in the statistical models because these variables were strongly correlated with the fraction of agricultural land use (**Figure S19**). Similarly, we did not use mean elevation in the statistical models because it showed a moderate correlation with air temperature (**Figure S19**). Correlations for other combinations of explanatory variables were weak to moderate.

Climate (air temperature and precipitation) and land use (the fraction of agricultural land use and dam density) differed clearly between the regions. Therefore, we used residuals of the simple linear regressions between the climate or land use variable (response variable) and region (explanatory variable). The residuals represent deviations from the regional averages; positive values indicate values greater than the regional average while negative values indicating values smaller than the regional average.

We constructed two models for each of the diversity metrics: global and region-specific models. The global model assumes that the effects of watershed area and branching probability are consistent across geographic regions (i.e., no interactive effects of the region with other linear predictors).

$$\mu_w = \xi_0 + \xi_1\ log_{10}\ WA_w + \xi_2\ log_{10}\ P_{b,w} + \xi_3\ region_w + \sum_k \xi_k x_{k,w} \tag{9}$$

where $\xi_k$ are the intercept and regression coefficients for linear predictors. Meanwhile, the region-specific model assumes the effects of watershed area and branching probability differ between regions (i.e., interactions between region and watershed area/branching probability).

$$\begin{aligned}\mu_w =& \xi_0 + \xi_1\ log_{10}\ WA_w + \xi_2\ log_{10}\ P_{b,w} + \xi_3\ region_w + \\ & \xi_4\ log_{10}\ WA_w \cdot region_w + \xi_5\ log_{10}\ P_{b,w} \cdot region_w + \sum_k \xi_k x_{k,w}\end{aligned} \tag{10}$$

All explanatory variables except watershed area, branching probability, and region were standardized to a mean of zero and a standard deviation of one before the analysis.

We compared the performance of these competing models using the Bayes factor (*35*), which gives the relative strength of evidence in favor of the global model (M0) over the region-specific model (M1). We used Bayesian Information Criteria (BIC) of the linear models to approximate the Bayes factor (*35*):

$$Bayes\ factor = exp(\Delta BIC_{10}/2) \tag{11}$$

where $\Delta BIC_{10} = BIC(M1) - BIC(M0)$.

For the supported model of $\gamma$ diversity, we estimated average predictive comparisons of watershed area and branching probability to compare their effect sizes. The average predictive comparison is an expected change of the response variable per a unit difference of the explanatory variable of interest (*36*). Unlike other common metrics of effect size (e.g., standardized regression coefficients), this method is applicable to models with variable transformations and allows direct comparisons of the effect sizes of watershed area and branching probability (see **Supplementary Text** for further details). We performed statistical analysis using R version 4.0.2 (*13*).

### Supplementary Text

#### Theory

##### Sensitivity analysis

We performed a sensitivity analysis of the metacommunity simulation to identify key simulation parameters that strongly affect the relationships between diversity metrics ($\alpha$, $\beta$, and $\gamma$) and ecosystem properties (the number of habitat patches $N_p$ and branching probability $P_b$). We generated 500 sets of parameter combinations by randomly drawing values of 8 simulation parameters from uniform distributions (**Table S1**). For each parameter combination, we generated 100 branching networks with the gradients of ecosystem size ($N_p \sim Unif(10, 150)$) and complexity ($P_b \sim Unif(0.01, 0.99)$). This results in a total of 50000 simulation replicates. In each simulation replicate, we allowed interspecific variation in niche optimum $\mu_i$ and width $\sigma_{niche,i}$ ($\mu_i \sim Unif(-1, 1)$ and $\sigma_{niche,i} \sim Unif(0.1, 1)$, respectively; subscript $i$ represents species) and ran 1400 time steps of metacommunity dynamics. We obtained temporal means of $\alpha$, $\beta$, and $\gamma$ diversity for the last 1000 time steps. The first 400 time steps were discarded as initialization and burn-in periods. We removed simulation replicates in which no species established populations over the initial 400 time steps.

For each parameter combination, we regressed log-transformed $\alpha$, $\beta$, and $\gamma$ diversity ($log_{10}\ y_j$ for network replicate $j$) on the number of habitat patches $N_p$ and branching probability $P_b$ as:

$$\begin{aligned} log_{10}\ y_j &\sim Normal(\mu_j, \sigma^2) \\ \mu_j &= \psi_0 + \psi_1\ log_{10}\ N_{p,j} + \psi_2\ log_{10}\ P_{b,j} \end{aligned} \quad (12)$$

where $\psi_k$ ($k = 0 - 2$) are the intercept ($\psi_0$) and regression coefficients ($\psi_1$ and $\psi_2$). We extracted 500 estimates of $\psi_1$ and $\psi_2$, which represent the effects of $N_p$ and $P_b$ on diversity metrics under a given parameter combination. To examine influences of simulation parameters (**Table S1**) on $\psi_1$ and $\psi_2$, we developed the following regression model taking $\psi_1$ or $\psi_2$ as a response variable $u_n$ (parameter combination $n$):

$$\begin{aligned} &u_n \sim Normal(\mu_n, \sigma^2) \\ &\mu_n = \zeta_0 + \zeta_1\sigma_{h,n} + \zeta_2\sigma_{l,n} + \zeta_3\sigma_{z,n} + \zeta_4\phi_n + \zeta_5\nu_n + \zeta_6\alpha_{max,n} + \zeta_7\theta_n + \zeta_8 p_{d,n} \end{aligned} \quad (13)$$

where $\zeta_k$ ($k = 0 - 8$) are the intercept ($\zeta_0$) and regression coefficients ($\zeta_{1-8}$). Explanatory variables were standardized to a mean of zero and a standard deviation of one, so that regression coefficients are comparable.

The sensitivity analysis revealed key simulation parameters. For the effects of $N_p$, the following simulation parameters were influential: the degree of local environmental noise ($\sigma_l$; influenced the effects on $\alpha$ and $\gamma$ diversity), the maximum value of interspecific competition coefficient ($\alpha_{max}$; influenced the effects on $\alpha$, $\beta$, and $\gamma$ diversity), dispersal distance ($\theta$; influenced the effects on $\alpha$ and $\beta$ diversity), and dispersal probability ($p_d$; influenced the effect on $\alpha$ diversity) (**Table S3**). For the effects of $P_b$, the following simulation parameters were influential: environmental variation at headwaters ($\sigma_h$; influenced the effect on $\gamma$ diversity), the degree of local environmental noise ($\sigma_l$; influenced the effects on $\alpha$ and $\beta$ diversity), and dispersal distance ($\theta$; influenced the effects on $\alpha$ and $\beta$ diversity) (**Table S4**).

Based on the results, we identified $\sigma_h$, $\sigma_l$, $\alpha_{max}$, $\theta$, and $p_d$ as key parameters. We changed the values of these parameters in the main analysis and examined the relationships between diversity metrics and ecosystem properties.

##### Longitudinal gradient of local species richness

Longitudinal gradients of local species richness have been extensively studied in rivers, illuminating typical patterns observed in nature. The most common pattern is a downstream increase of local species richness (*10*, *11*, *37*). However, recent empirical and theoretical studies also showed 'reversed patterns,' in which local species richness decreases downstream (*38*, *39*). We predicted the longitudinal gradient of local species richness to confirm that our simulation scenarios are capable of reproducing the previously observed patterns of local species richness. We considered 32 simulation scenarios comprising four landscape and eight ecological scenarios, as described in the main text (a set of parameters is described in **Table S2**). Under each simulation scenario, we generated 10 branching networks with fixed parameters of ecosystem size ($N_p = 100$) and complexity ($P_b = 0.5$). This results in a total of 320 simulation replicates. In each simulation replicate, we allowed interspecific variation in niche optimum $\mu_i$ and width $\sigma_{niche,i}$ ($\mu_i \sim Unif(-1, 1)$ and $\sigma_{niche,i} \sim Unif(0.1, 1)$, respectively; subscript $i$ represents species) and ran 1400 time steps of metacommunity dynamics. We obtained temporal means of local species richness at each habitat patch for the last 1000 time steps. The first 400 time steps were discarded as initialization and burn-in periods. We removed simulation replicates in which no species established populations over the initial 400 time steps. We evaluated the relationship between local species richness and the number of upstream habitat patches, a proxy for the longitudinal position of a habitat patch.

The simulation reproduced diverse patterns of longitudinal gradients in local species richness (**Figures S1-4**). The downstream increase of local species richness was predicted under a natural landscape scenario, in which environmental variation at headwaters $\sigma_h$ exceeds the degree of local environmental noise $\sigma_l$ (**Figure S1**). This pattern was consistent across ecological scenarios except those with long dispersal distance and high dispersal probability (**Figure S1**). Similarly, we observed a downstream increase of local species richness in scenarios with low habitat diversity ($\sigma_h = \sigma_l = 0.01$) and low dispersal probability ($p_d = 0.01$) (**Figure S3**). However, there were cases where local species richness decreased downstream or showed no longitudinal patterns. For example, when local environmental noise exceeds environmental variation at headwaters ($\sigma_l \geq \sigma_h$), local species richness showed a downstream decrease or a vague longitudinal pattern (**Figure S4**). Therefore, the simulation scenarios were capable of reproducing previously observed patterns, suggesting the appropriateness in the choice of parameter combinations.

#### Empirical data

##### Data selection criteria

**Hokkaido, Japan.** In Hokkaido, most data were collected from summer to fall. We screened data through the following procedure:

1. We listed recorded fish species and re-organized species names to make them consistent across data sources. We removed the following species at this stage: (1) identified at the family-level; (2) marine fish species (including species that occasionally use brackish/freshwater habitats).
2. We selected sampling sites based on the following criteria: (1) surveys were conducted with netting and/or electrofishing, (2) surveys were designed to collect a whole fish community, (3) sites contained reliable coordinates (sites with coordinates identical at 3 decimal degrees were treated as the same site), and (4) sites did not involve unidentified species (genus level) that are rarely observed in the data set (< 100 sites occurrence).
3. For sites with multiple visits (i.e., temporal replicates), we used the latest-year observation at each sampling site to minimize variation in sampling efforts among sites. Surveys that occurred in the same year were aggregated into a single observation.
4. We confined sites to those with the latest observation year of ≥ 1990. Although the data set contained observations from 1953, we added this restriction to align the observation period with the data set in the Midwest, US.
5. Four genera (*Lethenteron*, *Pungitius*, *Rhinogobius*, and *Tribolodon*) were treated as species groups (i.e., spp.) as their taxonomic resolutions varied greatly among data sources due to difficulties in identifying species.

**Midwest, US.** In the Midwest, the data set covered most of Upper Mississippi (Hydrologic Unit Code 2 [HUC 2] , region 07, as defined by U.S. Geological Survey and U.S. Department of Agriculture Natural Resources Conservation Service (*40*)) and the part of Great Lakes (HUC 2, region 04), Missouri (HUC 2, region 10), and Ohio (HUC 2, region 05). Fish data were collected from summer to fall with electrofishing (backpack, barge-type, or boat-mounted) and supplemental netting at some locations. We screened data through the following procedure:

1. We used data of the Upper Mississippi (HUC 2, region 07) and Great Lakes basins (HUC 2, region 04) as most sites are included in these regions.
2. We removed records of unidentified species, hybrid species, and commercial species apparently absent in the wild (e.g., goldfish).
3. We used the latest observation at each sampling site to minimize variation in sampling efforts among sites.

##### Asymptotic species richness

We evaluated sensitivity of asymptotic species richness (Chao 2 estimator) to sample size (i.e., the number of sampling sites in a watershed). We simulated presence-absence data of species with known values of true species richness $S_{true}$ and the number of sampling sites $N_{site}$. In this simulation, presence of species $i$ at site $x$, $J_{ix}$, was drawn from a Bernoulli distribution as $J_{ix} \sim Bernoulli(p_i \kappa_{ix})$ where $p_i$ is the detection probability for species $i$ and $\kappa_{ix}$ is the presence probability of species $i$ at site $x$. Based on the incidence frequency of simulated species $F_i = \sum_{x=1}^{N_{site}} J_{ix}$, we estimated asymptotic species richness using the iNEXT function in the R package 'iNEXT' (*23*). We calculated % bias of estimated asymptotic species richness $S_{est}$:

$$\% \, bias = \frac{100(S_{est} - S_{true})}{S_{true}} \tag{14}$$

Positive values of % bias indicate an overestimation of species richness, while negative values indicate an underestimation.

We used the following values for parameters: $S_{true} = 10, 40, 70, 100$ and $N_{site} = 5, 10, 15, 20, 100$. We produced 100 replicates of simulated data sets for each parameter combination, resulting in a total of 2000 simulation replicates. In each simulation replicate, the probabilities of detection and true presence were drawn randomly from uniform distributions as $p_i \sim Unif(0.3, 0.8)$ and $\kappa_{ix} \sim Unif(0, 1)$.

The number of sampling sites influenced the estimation accuracy of asymptotic species richness. The % bias decreased sharply as the number of sampling sites increased (**Figure S20**). In particular, the estimation bias with a small sample size ($N_{site} = 5$) was substantial when the true species richness $S_{true}$ was low; some estimates showed $> 150\%$ bias (**Figure S20**). Given the simulation results, estimates of asymptotic species richness at watersheds with $< 10$ sampling sites may involve substantial statistical uncertainty.

**Average predictive comparison**

For regression models, standardized regression coefficients (or its variant) are perhaps the most common summary when comparing effect sizes of explanatory variables. These values are useful if they are directly interpretable. For example, in linear regression models without interactions and variable transformations, regression coefficients have direct interpretations as they represent additive effects. However, there are many cases where regression coefficients are difficult to interpret. In our case, we regressed the log-transformed species richness on explanatory variables, so regression coefficients do not have direct interpretations at the original scale of the response variable $y$ (i.e., species richness).

The average predictive comparison provides an intuitive yet rigorous way to quantify effect sizes for each of explanatory variables in regression models with interactions and/or variable transformations (*36*). The basic predictive comparison $\delta_u$ is an expected change in $y$ on the original scale per a unit difference of the explanatory variable of interest:

$$\delta_u(u^{(1)} \rightarrow u^{(2)}, v, \Theta) = \frac{E(y|u^{(2)}, v, \Theta) - E(y|u^{(1)}, v, \Theta)}{u^{(2)} - u^{(1)}} \tag{15}$$

where $E(\cdot)$ is a known function (e.g., exponential), $u$ the input of interest (a value for the explanatory variable of interest), $v$ all the other inputs (a vector in general), $\Theta$ a set of parameters in a regression model. In general, $\delta_u$ rests on $u^{(1)}$ and $u^{(2)}$ (the lower and higher points of the hypothesized change in the explanatory variable of interest), $v$, and $\Theta$. Therefore, Gelman and Pardoe (*36*) defined the *average predictive comparison* $\Delta_u$, the mean value of $\delta_u$ over specified distributions of the inputs and model parameters:

$$\Delta_u = \frac{\int\int_{u^{(2)}>u^{(1)}} du^{(1)}du^{(2)} \int dv \int d\Theta (E(y|u^{(2)}, v, \Theta) - E(y|u^{(1)}, v, \Theta)) \; p(u^{(1)}|v) \; p(u^{(2)}|v) \; p(v) \; p(\Theta)}{\int\int_{u^{(2)}>u^{(1)}} du^{(1)}du^{(2)} \int dv \int d\Theta \; (u^{(2)} - u^{(1)}) \; p(u^{(1)}|v) \; p(u^{(2)}|v) \; p(v) \; p(\Theta)} \tag{16}$$

However, directly using the above equation is impractical because estimating $p(u^{(1)}|v)$ and $p(u^{(2)}|v)$ from finite data points is challenging (especially when $v$ is continuous). Instead, we estimated $\Delta_u$ using the following equation proposed by Gelman and Pardoe (*36*):

$$\hat{\Delta}_u = \frac{\sum_{i=1}^{n}\sum_{j=1}^{n}\sum_{s=1}^{S} w_{ij}(E(y|u_j, v_i, \Theta_s) - E(y|u_i, v_i, \Theta_s))sign(u_j - u_i)}{\sum_{i=1}^{n}\sum_{j=1}^{n}\sum_{s=1}^{S} w_{ij}(u_j - u_i)sign(u_j - u_i)} \tag{17}$$

The summations over $n$ data ($i$ and $j$ are a given data point) and $S$ parameter replicates ($s$ is a given simulation replicate) are a realization of averaging over the distributions of $(u^{(1)}, v)$ , $u^{(2)}$, and $\Theta$. The factor $w_{ij}$ is a weight that serves to approximate $p(u^{(1)}|v)$ and $p(u^{(2)}|v)$. In theory, $v$ must be held constant while the input of interest changes from $u^{(1)}$ to $u^{(2)}$. However, there are, if any, few transitions from $u^{(1)}$ to $u^{(2)}$ with a common $v$. We approximate such exact transitions by assigning each pair of data points with a weight:

$$w_{ij} = w(v_i, v_j) \tag{18}$$

The weight factor should represent the likelihood of $u$ changing from $u_i$ to $u_j$ with a common $v = v_i$ (note that $v_i$ is in general a vector of explanatory variables for data point $i$). We used the following weighting function based on Mahalanobis distances following Gelman and Pardoe (*36*):

$$w(v_i, v_j) = \frac{1}{1 + (v_i - v_j)^T \Omega_v^{-1} (v_i - v_j)} \quad (19)$$

where $\Omega_v^{-1}$ is the inverse of the variance-covariance matrix of $v$. The function gives the maximum weight for a pair of $u_i$ and $u_j$ if $v_i = v_j$ (note that $v = v_i$ when $u = u_i$) while giving a less weight as the Mahalanobis distance between $v_i$ and $v_j$ increases. This property makes sense because our goal is to approximate the probability of transition from $u_i$ to $u_j$ with a common $v$ (i.e., $v_i = v_j$).

We estimated $\hat{\Delta}_u$ for watershed area and branching probability using the estimated coefficients $\hat{\xi}_k$ of the regression model explaining $\gamma$ diversity. A thousand of vectors of simulated parameters $\Theta_s$ were drawn from normal distributions with means of $\hat{\xi}_k$ and standard deviations of $\sigma_{\xi,k}$ (the estimated standard errors of parameters $\xi_k$). The estimated average predictive comparisons are $\hat{\Delta}_u = \frac{1}{S}\sum_{s=1}^{S} \hat{\Delta}_{u,s}$ where

$$\hat{\Delta}_{u,s} = \frac{\sum_{i=1}^{n}\sum_{j=1}^{n} w_{ij}(E(y|u_j, v_i, \Theta_s) - E(y|u_i, v_i, \Theta_s))sign(u_j - u_i)}{\sum_{i=1}^{n}\sum_{j=1}^{n} w_{ij}(10^{u_j} - 10^{u_i})sign(u_j - u_i)} \quad (20)$$

We exponentiated $u_i$ and $u_j$ in the denominator because we log-transformed watershed area and branching probability in the statistical model. Then

$$S.E.(\hat{\Delta}_{u,s}) = \sqrt{\frac{1}{S-1}\sum_{s=1}^{S}(\hat{\Delta}_{u,s} - \hat{\Delta}_u)^2} \quad (21)$$

The estimated average predictive comparisons were summarized in **Table S7**.

### Tables

##### Table S1 Simulation parameter (sensitivity analysis)

**Table S1** Parameter values used in the sensitivity analysis of the metacommunity simulation. See the main text for model details.

| Parameter | Value | Interpretation |
|---|---|---|
| $\sigma_h$ | Unif(0.01, 1) | Environmental variation at headwaters |
| $\sigma_l$ | Unif(0.01, 1) | Degree of local environmental noise |
| $\sigma_z$ | Unif(0.01, 0.5) | Temporal environmental variability |
| $\rho$ | 1 | Strength of spatial autocorrelation in mean environmental condition |
| $\phi$ | Unif(0.01, 1) | Extent of spatial autocorrelation in temporal environmental variation |
| $\nu$ | Unif(1, 5) | Cost of a wider niche |
| $\alpha_{max}$ | Unif(0.5, 1.5) | Maximum value of interspecific competition coefficient |
| $\theta$ | Unif(0.1, 1) | Rate parameter of an exponential dispersal kernel |
| $p_d$ | Unif(0.01, 0.1) | Dispersal probability |
| $r_{0,i}$ | 4 | Maximum reproductive rate |

#### Table S2 Simulation parameter (main analysis)

**Table S2** Values and interpretation of simulation parameters used in the main simulation. See the main text for model details.

| Parameter | Value | Interpretation |
|---|---|---|
| $\sigma_h$ | 0.01, 1.00 | Environmental variation at headwaters |
| $\sigma_l$ | 0.01, 1.00 | Degree of local environmental noise |
| $\sigma_z$ | 0.1 | Temporal environmental variability |
| $\rho$ | 1 | Strength of spatial autocorrelation in mean environmental condition |
| $\phi$ | 0.05 | Extent of spatial autocorrelation in temporal environmental variation |
| $\nu$ | 1 | Cost of a wider niche |
| $\alpha_{max}$ | 0.75, 1.50 | Maximum value of interspecific competition coefficient |
| $\theta$ | 0.1, 1.0 | Rate parameter of an exponential dispersal kernel |
| $p_d$ | 0.01, 0.10 | Dispersal probability |
| $r_{0,i}$ | 4 | Maxiumum reproductive rate |

**Table S3 Sensitivity analysis for ecosystem size effects**

**Table S3** Sensitivity analysis of the metacommunity simulation. Parameter estimates of linear regression models (standard errors in parenthesis) are shown. Response variables are the effects of the number of habitat patches ($N_p$) on $\alpha$, $\beta$, and $\gamma$ diversity. Explanatory variables (i.e., simulation parameters) were standardized to a mean of zero and a standard deviation of one prior to the analysis. See Tables S1 and S2 for interpretation of the simulation parameters.

| | Response variable | | |
|---|---|---|---|
| | Effect of $N_p$ on $\alpha$ diversity | Effect of $N_p$ on $\beta$ diversity | Effect of $N_p$ on $\gamma$ diversity |
| $\sigma_h$ | 0.008 | −0.003 | 0.005 |
| | (0.002) | (0.001) | (0.002) |
| $\sigma_l$ | −0.021 | 0.004 | −0.018 |
| | (0.002) | (0.001) | (0.002) |
| $\sigma_z$ | 0.0001 | −0.013 | −0.013 |
| | (0.002) | (0.001) | (0.002) |
| $\phi$ | 0.001 | −0.0002 | 0.0003 |
| | (0.002) | (0.001) | (0.002) |
| $\nu$ | −0.001 | −0.009 | −0.009 |
| | (0.002) | (0.001) | (0.002) |
| $\alpha_{max}$ | 0.019 | 0.028 | 0.047 |
| | (0.002) | (0.001) | (0.002) |
| $\theta$ | −0.040 | 0.041 | 0.001 |
| | (0.002) | (0.001) | (0.002) |
| $p_d$ | 0.017 | −0.006 | 0.010 |
| | (0.002) | (0.001) | (0.002) |
| Intercept | 0.147 | 0.137 | 0.284 |
| | (0.002) | (0.001) | (0.002) |

**Table S4 Sensitivity analysis for ecosystem complexity effects**

**Table S4** Sensitivity analysis of the metacommunity simulation. Parameter estimates of linear regression models (standard errors in parenthesis) are shown. Response variables are the effects of branching probability ($P_b$) on $\alpha$, $\beta$, and $\gamma$ diversity. Explanatory variables (i.e., simulation parameters) were standardized to a mean of zero and a standard deviation of one prior to the analysis. See Tables S1 and S2 for interpretation of the simulation parameters.

| | Response variable | | |
|---|---|---|---|
| | Effect of $P_b$ on $\alpha$ diversity | Effect of $P_b$ on $\beta$ diversity | Effect of $P_b$ on $\gamma$ diversity |
| $\sigma_h$ | 0.012<br>(0.003) | 0.007<br>(0.002) | 0.019<br>(0.002) |
| $\sigma_l$ | 0.060<br>(0.003) | −0.047<br>(0.002) | 0.013<br>(0.002) |
| $\sigma_z$ | −0.004<br>(0.003) | −0.002<br>(0.002) | −0.006<br>(0.002) |
| $\phi$ | 0.002<br>(0.003) | 0.001<br>(0.002) | 0.002<br>(0.002) |
| $\nu$ | −0.001<br>(0.003) | 0.001<br>(0.002) | −0.001<br>(0.002) |
| $\alpha_{max}$ | −0.006<br>(0.003) | −0.001<br>(0.002) | −0.006<br>(0.002) |
| $\theta$ | 0.027<br>(0.003) | −0.028<br>(0.002) | −0.001<br>(0.002) |
| $p_d$ | 0.007<br>(0.003) | −0.007<br>(0.002) | −0.0002<br>(0.002) |
| Intercept | 0.145<br>(0.003) | −0.132<br>(0.002) | 0.013<br>(0.002) |

**Table S5 List of fish species in Hokkaido, Japan**

**Table S5** List of fish species in Hokkaido, Japan, included in our statistical analysis. 0 species are ordered alphabetically, along with the number of sites present and % occupancy out of 0 sites.

| Species | Number of sites present | Occupancy (%) |
|---|---|---|
| Acanthogobius lactipes | 65 | 2.49 |
| Anguilla japonica | 1 | 0.04 |
| Carassius buergeri subsp. 2 | 4 | 0.15 |
| Carassius cuvieri | 24 | 0.92 |
| Carassius sp. | 213 | 8.17 |
| Channa argus | 3 | 0.12 |
| Cottus amblystomopsis | 49 | 1.88 |
| Cottus hangiongensis | 94 | 3.61 |
| Cottus nozawae | 840 | 32.22 |
| Cottus sp. ME | 25 | 0.96 |
| Cyprinus carpio | 50 | 1.92 |
| Gasterosteus aculeatus | 151 | 5.79 |
| Gnathopogon caerulescens | 1 | 0.04 |
| Gnathopogon elongatus elongatus | 2 | 0.08 |
| Gymnogobius breunigii | 29 | 1.11 |
| Gymnogobius castaneus complex | 145 | 5.56 |
| Gymnogobius opperiens | 87 | 3.34 |
| Gymnogobius petschiliensis | 2 | 0.08 |
| Gymnogobius urotaenia | 293 | 11.24 |
| Hucho perryi | 61 | 2.34 |
| Hypomesus nipponensis | 171 | 6.56 |
| Hypomesus olidus | 8 | 0.31 |
| Lefua nikkonis | 20 | 0.77 |
| Lethenteron spp. | 739 | 28.35 |
| Leucopsarion petersii | 3 | 0.12 |
| Luciogobius guttatus | 3 | 0.12 |
| Misgurnus anguillicaudatus | 215 | 8.25 |
| Noemacheilus barbatulus | 1601 | 61.41 |
| Oncorhynchus gorbuscha | 27 | 1.04 |
| Oncorhynchus keta | 149 | 5.72 |
| Oncorhynchus masou masou | 1418 | 54.39 |
| Oncorhynchus mykiss | 464 | 17.80 |
| Oncorhynchus nerka | 6 | 0.23 |
| Opsariichthys platypus | 1 | 0.04 |
| Osmerus dentex | 6 | 0.23 |
| Phoxinus percnurus sachalinensis | 68 | 2.61 |
| Plecoglossus altivelis altivelis | 111 | 4.26 |
| Pseudorasbora parva | 93 | 3.57 |
| Pungitius spp. | 286 | 10.97 |
| Rhinogobius spp. | 175 | 6.71 |
| Rhodeus ocellatus ocellatus | 22 | 0.84 |
| Salangichthys microdon | 11 | 0.42 |
| Salmo trutta | 15 | 0.58 |
| Salvelinus fontinalis | 2 | 0.08 |
| Salvelinus leucomaenis leucomaenis | 625 | 23.97 |
| Salvelinus malma | 274 | 10.51 |
| Salvelinus malma miyabei | 2 | 0.08 |
| Silurus asotus | 7 | 0.27 |

| Species | Number of sites present | Occupancy (%) |
|---|---:|---:|
| Spirinchus lanceolatus | 7 | 0.27 |
| Tribolodon spp. | 1171 | 44.92 |
| Tridentiger brevispinis | 137 | 5.26 |
| Tridentiger obscurus | 7 | 0.27 |

**Table S6 List of fish species in Midwest, US**

**Table S6** List of fish species in the Midwest, US, included in our statistical analysis. 0 species are ordered alphabetically, along with the number of sites present and % occupancy out of 0 sites.

| Species | Number of sites present | Occupancy (%) |
|---|---:|---:|
| Acipenser fulvescens | 7 | 0.18 |
| Alosa pseudoharengus | 1 | 0.03 |
| Ambloplites rupestris | 707 | 17.68 |
| Ameiurus melas | 868 | 21.71 |
| Ameiurus natalis | 665 | 16.63 |
| Ameiurus nebulosus | 30 | 0.75 |
| Amia calva | 95 | 2.38 |
| Ammocrypta clara | 12 | 0.30 |
| Aphredoderus sayanus | 76 | 1.90 |
| Aplodinotus grunniens | 208 | 5.20 |
| Campostoma anomalum | 1347 | 33.69 |
| Campostoma oligolepis | 125 | 3.13 |
| Carpiodes carpio | 128 | 3.20 |
| Carpiodes cyprinus | 234 | 5.85 |
| Carpiodes velifer | 82 | 2.05 |
| Catostomus commersonii | 2931 | 73.31 |
| Centrarchus macropterus | 5 | 0.13 |
| Chrosomus eos | 336 | 8.40 |
| Chrosomus neogaeus | 103 | 2.58 |
| Clinostomus elongatus | 97 | 2.43 |
| Cottus bairdii | 467 | 11.68 |
| Cottus carolinae | 6 | 0.15 |
| Cottus cognatus | 38 | 0.95 |
| Crystallaria asprella | 1 | 0.03 |
| Ctenopharyngodon idella | 19 | 0.48 |
| Culaea inconstans | 1531 | 38.29 |
| Cyprinella lutrensis | 269 | 6.73 |
| Cyprinella spiloptera | 780 | 19.51 |
| Cyprinella venusta | 2 | 0.05 |
| Cyprinella whipplei | 33 | 0.83 |
| Cyprinus carpio | 946 | 23.66 |
| Dorosoma cepedianum | 208 | 5.20 |
| Erimystax x-punctatus | 13 | 0.33 |
| Erimyzon oblongus | 63 | 1.58 |
| Erimyzon sucetta | 10 | 0.25 |
| Esox americanus vermiculatus | 117 | 2.93 |
| Esox lucius | 957 | 23.94 |
| Esox masquinongy | 20 | 0.50 |
| Etheostoma asprigene | 10 | 0.25 |
| Etheostoma blennioides | 1 | 0.03 |
| Etheostoma caeruleum | 195 | 4.88 |
| Etheostoma chlorosomum | 4 | 0.10 |
| Etheostoma crossopterum | 4 | 0.10 |
| Etheostoma exile | 261 | 6.53 |
| Etheostoma flabellare | 843 | 21.09 |
| Etheostoma gracile | 15 | 0.38 |
| Etheostoma kennicotti | 1 | 0.03 |
| Etheostoma microperca | 20 | 0.50 |

| Species | Number of sites present | Occupancy (%) |
|---|---|---|
| Etheostoma nigrum | 2546 | 63.68 |
| Etheostoma proeliare | 2 | 0.05 |
| Etheostoma spectabile | 121 | 3.03 |
| Etheostoma squamiceps | 4 | 0.10 |
| Etheostoma zonale | 321 | 8.03 |
| Fundulus diaphanus | 3 | 0.08 |
| Fundulus dispar | 4 | 0.10 |
| Fundulus notatus | 282 | 7.05 |
| Fundulus olivaceus | 42 | 1.05 |
| Hiodon alosoides | 3 | 0.08 |
| Hiodon tergisus | 16 | 0.40 |
| Hybognathus hankinsoni | 576 | 14.41 |
| Hybognathus nuchalis | 24 | 0.60 |
| Hybopsis amnis | 1 | 0.03 |
| Hypentelium nigricans | 682 | 17.06 |
| Hypophthalmichthys molitrix | 14 | 0.35 |
| Hypophthalmichthys nobilis | 3 | 0.08 |
| Ichthyomyzon castaneus | 51 | 1.28 |
| Ichthyomyzon fossor | 35 | 0.88 |
| Ichthyomyzon gagei | 6 | 0.15 |
| Ichthyomyzon unicuspis | 9 | 0.23 |
| Ictalurus punctatus | 413 | 10.33 |
| Ictiobus bubalus | 90 | 2.25 |
| Ictiobus cyprinellus | 129 | 3.23 |
| Ictiobus niger | 41 | 1.03 |
| Labidesthes sicculus | 64 | 1.60 |
| Lepisosteus oculatus | 13 | 0.33 |
| Lepisosteus osseus | 25 | 0.63 |
| Lepisosteus platostomus | 58 | 1.45 |
| Lepomis cyanellus | 1575 | 39.39 |
| Lepomis gibbosus | 290 | 7.25 |
| Lepomis gulosis | 43 | 1.08 |
| Lepomis humilis | 356 | 8.90 |
| Lepomis macrochirus | 1051 | 26.29 |
| Lepomis megalotis | 186 | 4.65 |
| Lepomis microlophus | 19 | 0.48 |
| Lethenteron appendix | 122 | 3.05 |
| Lota lota | 266 | 6.65 |
| Luxilus chrysocephalus | 198 | 4.95 |
| Luxilus cornutus | 1784 | 44.62 |
| Lythrurus fumeus | 6 | 0.15 |
| Lythrurus umbratilis | 224 | 5.60 |
| Macrhybopsis aestivalis | 1 | 0.03 |
| Macrhybopsis hyostoma | 3 | 0.08 |
| Macrhybopsis storeriana | 10 | 0.25 |
| Micropterus dolomieu | 748 | 18.71 |
| Micropterus punctulatus | 15 | 0.38 |
| Micropterus salmoides | 987 | 24.69 |
| Minytrema melanops | 41 | 1.03 |
| Morone americana | 2 | 0.05 |
| Morone chrysops | 65 | 1.63 |
| Morone mississippiensis | 16 | 0.40 |

| Species | Number of sites present | Occupancy (%) |
|---|---|---|
| Moxostoma anisurum | 288 | 7.20 |
| Moxostoma carinatum | 8 | 0.20 |
| Moxostoma duquesni | 103 | 2.58 |
| Moxostoma erythrurum | 709 | 17.73 |
| Moxostoma macrolepidotum | 737 | 18.43 |
| Moxostoma valenciennesi | 85 | 2.13 |
| Neogobius melanostomus | 13 | 0.33 |
| Nocomis biguttatus | 1287 | 32.19 |
| Notemigonus crysoleucas | 377 | 9.43 |
| Notropis anogenus | 10 | 0.25 |
| Notropis atherinoides | 207 | 5.18 |
| Notropis blennius | 16 | 0.40 |
| Notropis boops | 9 | 0.23 |
| Notropis buccatus | 47 | 1.18 |
| Notropis chalybaeus | 11 | 0.28 |
| Notropis dorsalis | 1088 | 27.21 |
| Notropis heterodon | 28 | 0.70 |
| Notropis heterolepis | 187 | 4.68 |
| Notropis hudsonius | 81 | 2.03 |
| Notropis nubilus | 58 | 1.45 |
| Notropis percobromus | 167 | 4.18 |
| Notropis rubellus | 63 | 1.58 |
| Notropis stramineus | 975 | 24.39 |
| Notropis texanus | 20 | 0.50 |
| Notropis volucellus | 81 | 2.03 |
| Notropis wickliffi | 10 | 0.25 |
| Noturus exilis | 34 | 0.85 |
| Noturus flavus | 479 | 11.98 |
| Noturus gyrinus | 447 | 11.18 |
| Noturus nocturnus | 38 | 0.95 |
| Oncorhynchus mykiss | 55 | 1.38 |
| Oncorhynchus tshawytscha | 1 | 0.03 |
| Opsopoeodus emiliae | 3 | 0.08 |
| Perca flavescens | 622 | 15.56 |
| Percina caprodes | 337 | 8.43 |
| Percina carprodes semifasciata | 7 | 0.18 |
| Percina evides | 16 | 0.40 |
| Percina maculata | 888 | 22.21 |
| Percina phoxocephala | 259 | 6.48 |
| Percina sciera | 2 | 0.05 |
| Percopsis omiscomaycus | 21 | 0.53 |
| Phenacobius mirabilis | 259 | 6.48 |
| Phoxinus erythrogaster | 417 | 10.43 |
| Pimephales notatus | 1784 | 44.62 |
| Pimephales promelas | 1535 | 38.39 |
| Pimephales vigilax | 84 | 2.10 |
| Pomoxis annularis | 61 | 1.53 |
| Pomoxis nigromaculatus | 376 | 9.40 |
| Pylodictis olivaris | 73 | 1.83 |
| Rhinichthys atratulus | 1120 | 28.01 |
| Rhinichthys cataractae | 627 | 15.68 |
| Rhinichthys obtusus | 449 | 11.23 |

| Species | Number of sites present | Occupancy (%) |
|---|---|---|
| Salmo trutta | 399 | 9.98 |
| Salvelinus fontinalis | 369 | 9.23 |
| Sander canadensis | 39 | 0.98 |
| Sander vitreus | 367 | 9.18 |
| Scaphirhynchus platorynchus | 8 | 0.20 |
| Semotilus atromaculatus | 2776 | 69.43 |
| Umbra limi | 1600 | 40.02 |

**Table S7 Average predictive comparisons**

Estimated average predictive comparisons. Regression coefficients were derived from the regression model explaining $\gamma$ diversity (see **Table 1** in the maintext for the estimated regression parameters). The average predictive comparisons (an expected increase in $\gamma$ diversity per a unit difference of the explanatory variable of interest) were estimated based on units of 1000 $km^2$ for watershed area and 0.1 for branching probability.

| Input | Estimate | SE | Unit |
|---|---|---|---|
| Watershed area | 5.75 | 3.18 | 1000 $km^2$ |
| Branching probability | 10.98 | 4.49 | 0.1 |

#### Figures

**Figure S1 Longitudinal gradient of local species richness ($\sigma_h = 1$, $\sigma_l = 0.01$)**

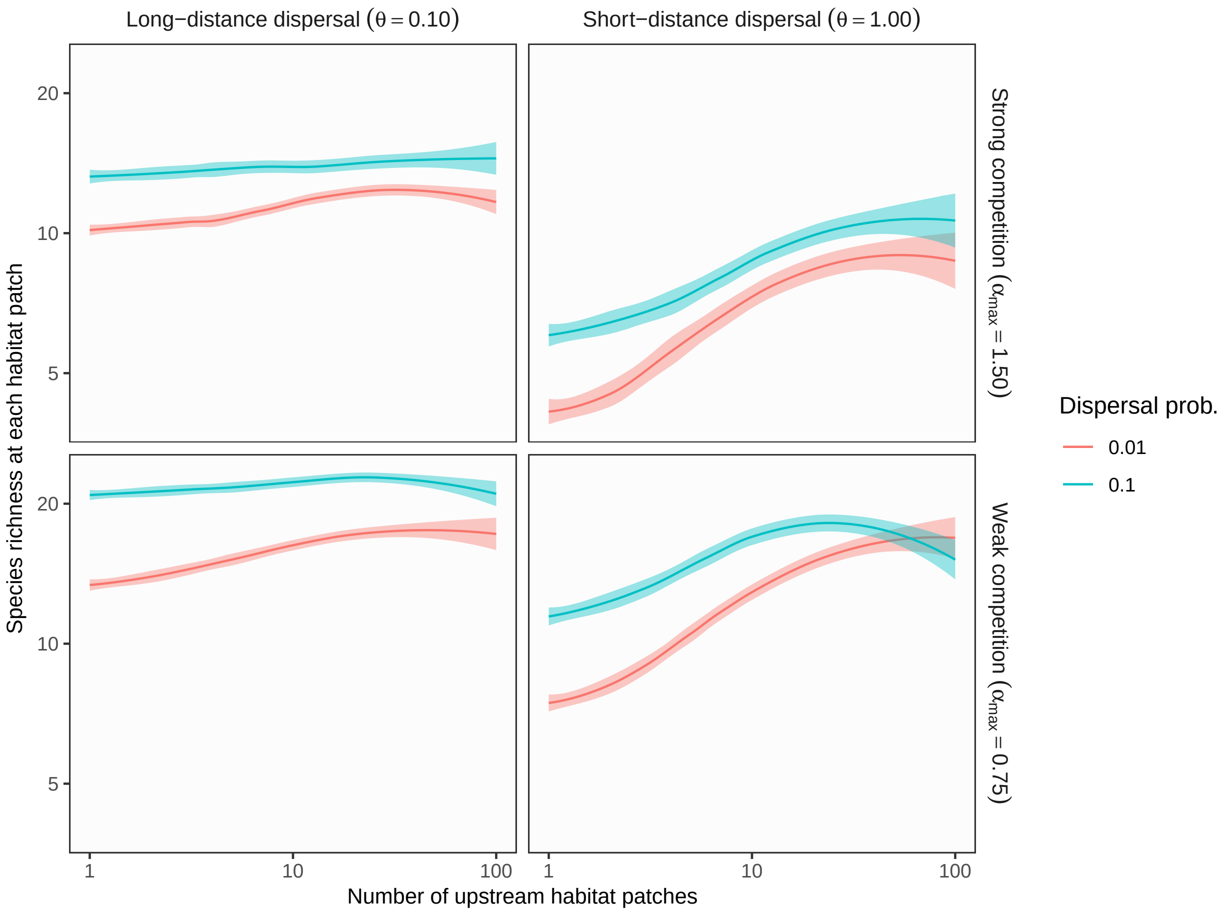

**Figure S1** Theoretical predictions for longitudinal gradients of local species richness in branching networks. The longitudinal position of each habitat patch (x-axis) was expressed as the number of upstream habitat patches. In this simulation, environmental variation at headwaters ($\sigma_h$) exceeds local environmental noise ($\sigma_l$). Lines and shades are loess curves fitted to simulated data and their 95% confidence intervals. Each panel represents different ecological scenarios under which metacommunity dynamics were simulated. Rows represent different competition strength. Competitive coefficients ($\alpha_{ij}$) were varied randomly from 0 to 1.5 (top, strong competition) or 0.75 (bottom, weak competition). Columns and lines represent different dispersal scenarios (dispersal distance and probability). Left and right columns show long-distance (the rate parameter of an exponential dispersal kernel $\theta = 0.10$) and short-distance dispersal ($\theta = 1.0$) scenarios repectively. Red and blue lines show low ($p_d = 0.01$) and high dispersal probabilities ($p_d = 0.10$). Other parameters are as follows: environmental variation at headwaters $\sigma_h = 1$; local environmental noise $\sigma_l = 0.01$; ecosystem size $N_p = 100$; ecosystem complexity $P_b = 0.5$.

**Figure S2 Longitudinal gradient of local species richness ($\sigma_h = 1$, $\sigma_l = 1$)**

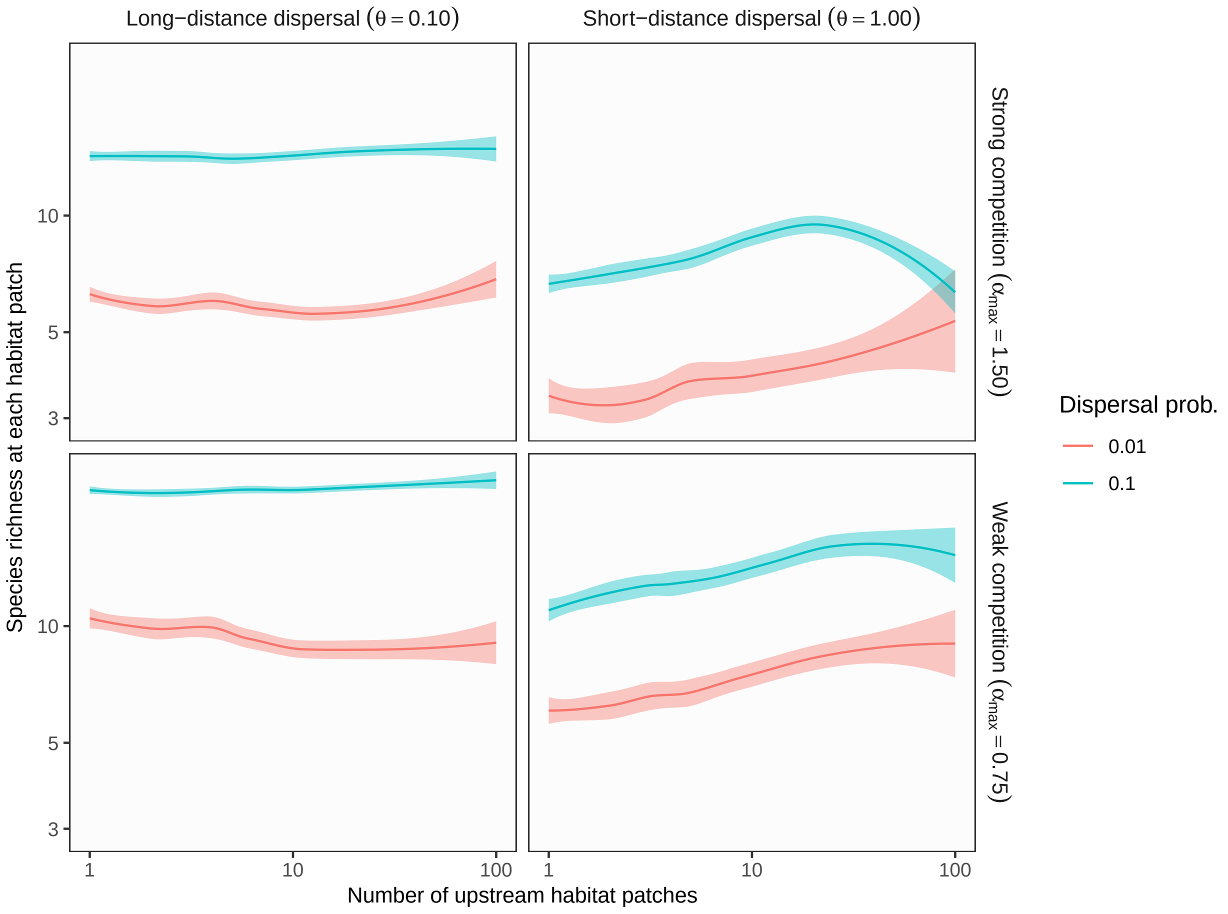

**Figure S2** Theoretical predictions for longitudinal gradients of local species richness in branching networks. The longitudinal position of each habitat patch (x-axis) was expressed as the number of upstream habitat patches. In this simulation, environmental variation at headwaters ($\sigma_h$) is equal to local environmental noise ($\sigma_l$). Lines and shades are loess curves fitted to simulated data and their 95% confidence intervals. Each panel represents different ecological scenarios under which metacommunity dynamics were simulated. Rows represent different competition strength. Competitive coefficients ($\alpha_{ij}$) were varied randomly from 0 to 1.5 (top, strong competition) or 0.75 (bottom, weak competition). Columns and lines represent different dispersal scenarios (dispersal distance and probability). Left and right columns show long-distance (the rate parameter of an exponential dispersal kernel $\theta = 0.10$) and short-distance dispersal ($\theta = 1.0$) scenarios repectively. Red and blue lines show low ($p_d = 0.01$) and high dispersal probabilities ($p_d = 0.10$). Other parameters are as follows: environmental variation at headwaters $\sigma_h = 1$; local environmental noise $\sigma_l = 1$; ecosystem size $N_p = 100$; ecosystem complexity $P_b = 0.5$.

**Figure S3 Longitudinal gradient of local species richness ($\sigma_h = 0.01$, $\sigma_l = 0.01$)**

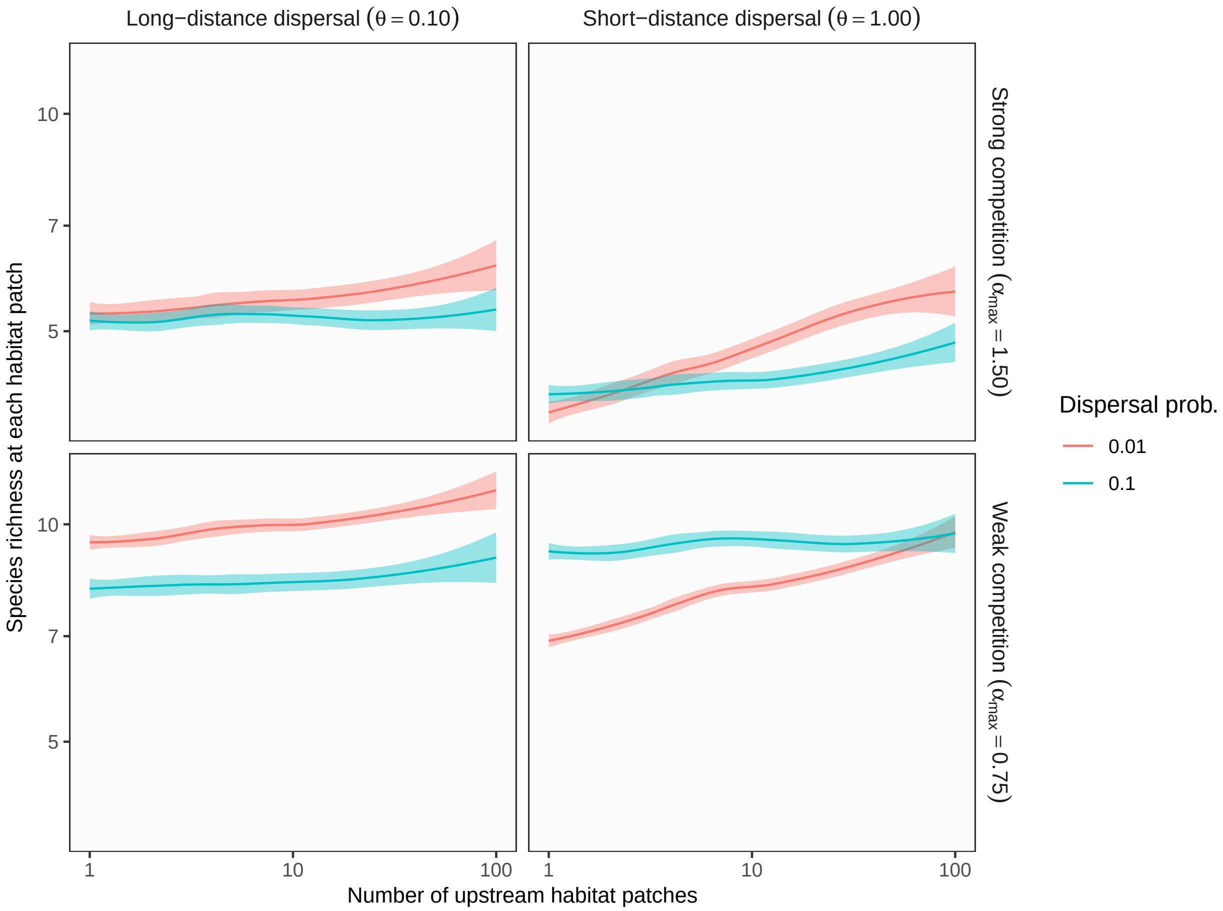

**Figure S3** Theoretical predictions for longitudinal gradients of local species richness in branching networks. The longitudinal position of each habitat patch (x-axis) was expressed as the number of upstream habitat patches. In this simulation, environmental variation at headwaters ($\sigma_h$) is equal to local environmental noise ($\sigma_l$). Lines and shades are loess curves fitted to simulated data and their 95% confidence intervals. Each panel represents different ecological scenarios under which metacommunity dynamics were simulated. Rows represent different competition strength. Competitive coefficients ($\alpha_{ij}$) were varied randomly from 0 to 1.5 (top, strong competition) or 0.75 (bottom, weak competition). Columns and lines represent different dispersal scenarios (dispersal distance and probability). Left and right columns show long-distance (the rate parameter of an exponential dispersal kernel $\theta = 0.10$) and short-distance dispersal ($\theta = 1.0$) scenarios repectively. Red and blue lines show low ($p_d = 0.01$) and high dispersal probabilities ($p_d = 0.10$). Other parameters are as follows: environmental variation at headwaters $\sigma_h = 0.01$; local environmental noise $\sigma_l = 0.01$; ecosystem size $N_p = 100$; ecosystem complexity $P_b = 0.5$.

**Figure S4 Longitudinal gradient of local species richness ($\sigma_h = 0.01$, $\sigma_l = 1$)**

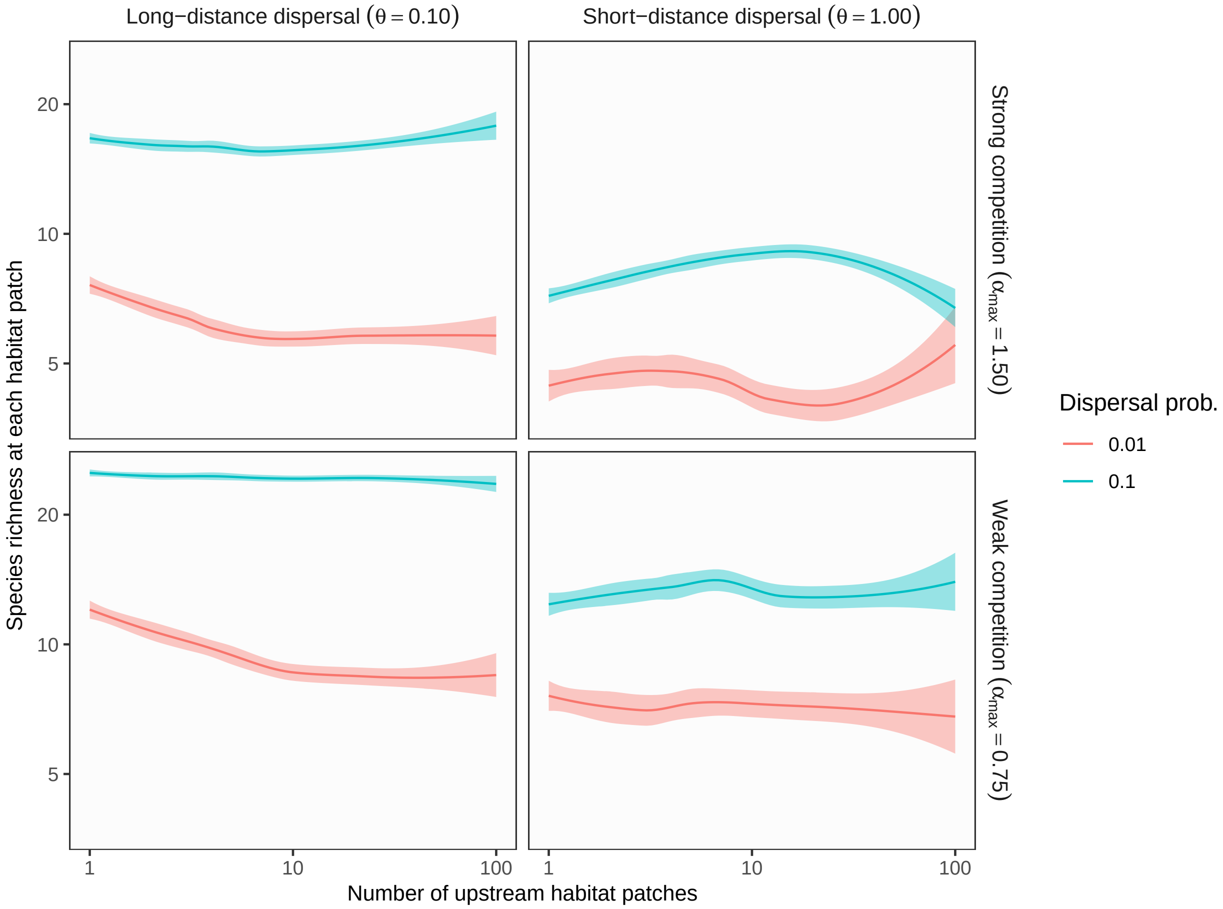

**Figure S4** Theoretical predictions for longitudinal gradients of local species richness in branching networks. The longitudinal position of each habitat patch (x-axis) was expressed as the number of upstream habitat patches. In this simulation, environmental variation at headwaters ($\sigma_h$) is less than local environmental noise ($\sigma_l$). Lines and shades are loess curves fitted to simulated data and their 95% confidence intervals. Each panel represents different ecological scenarios under which metacommunity dynamics were simulated. Rows represent different competition strength. Competitive coefficients ($\alpha_{ij}$) were varied randomly from 0 to 1.5 (top, strong competition) or 0.75 (bottom, weak competition). Columns and lines represent different dispersal scenarios (dispersal distance and probability). Left and right columns show long-distance (the rate parameter of an exponential dispersal kernel $\theta = 0.10$) and short-distance dispersal ($\theta = 1.0$) scenarios repectively. Red and blue lines show low ($p_d = 0.01$) and high dispersal probabilities ($p_d = 0.10$). Other parameters are as follows: environmental variation at headwaters $\sigma_h = 0.01$; local environmental noise $\sigma_l = 1$; ecosystem size $N_p = 100$; ecosystem complexity $P_b = 0.5$.

**Figure S5 Influence of ecosystem size** ($p_d = 0.1$, $\sigma_h = 1$, $\sigma_l = 0.01$)

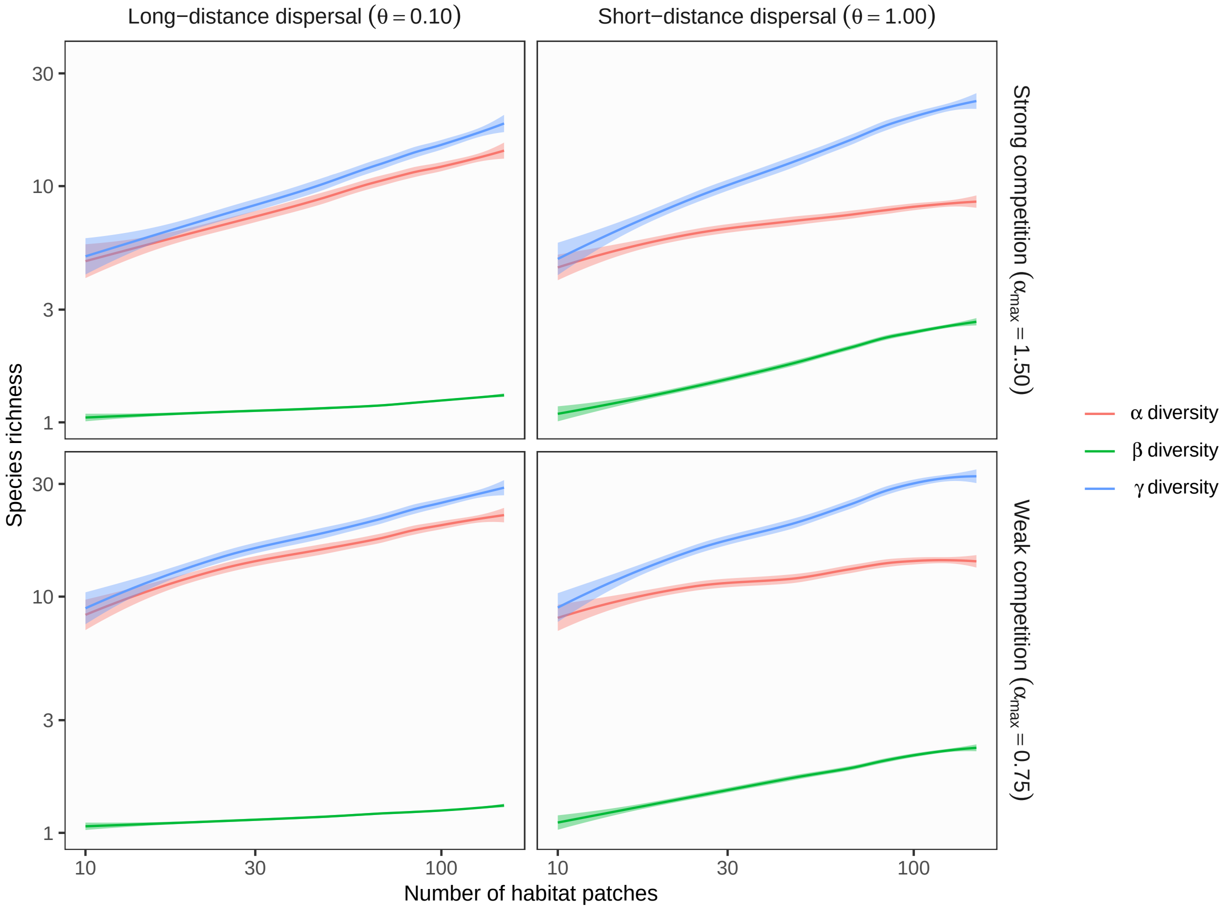

**Figure S5** Theoretical predictions for ecosystem size influences (the number of habitat patches) on $\alpha$, $\beta$, and $\gamma$ diversity in branching networks. In this simulation, environmental variation at headwaters ($\sigma_h$) exceeds local environmental noise ($\sigma_l$). Lines and shades are loess curves fitted to simulated data and their 95% confidence intervals. Each panel represents different ecological scenarios under which metacommunity dynamics were simulated. Rows represent different competition strength. Competitive coefficients ($\alpha_{ij}$) were varied randomly from 0 to 1.5 (top, strong competition) or 0.75 (bottom, weak competition). Columns represent different dispersal scenarios. Two dispersal parameters were chosen to simulate scenarios with long-distance (the rate parameter of an exponential dispersal kernel $\theta = 0.10$) and short-distance dispersal ($\theta = 1.0$). Other parameters are as follows: dispersal probability $p_d = 0.1$; environmental variation at headwaters $\sigma_h = 1$; local environmental noise $\sigma_l = 0.01$.

**Figure S6 Influence of ecosystem size** ($p_d = 0.1$, $\sigma_h = 1$, $\sigma_l = 1$)

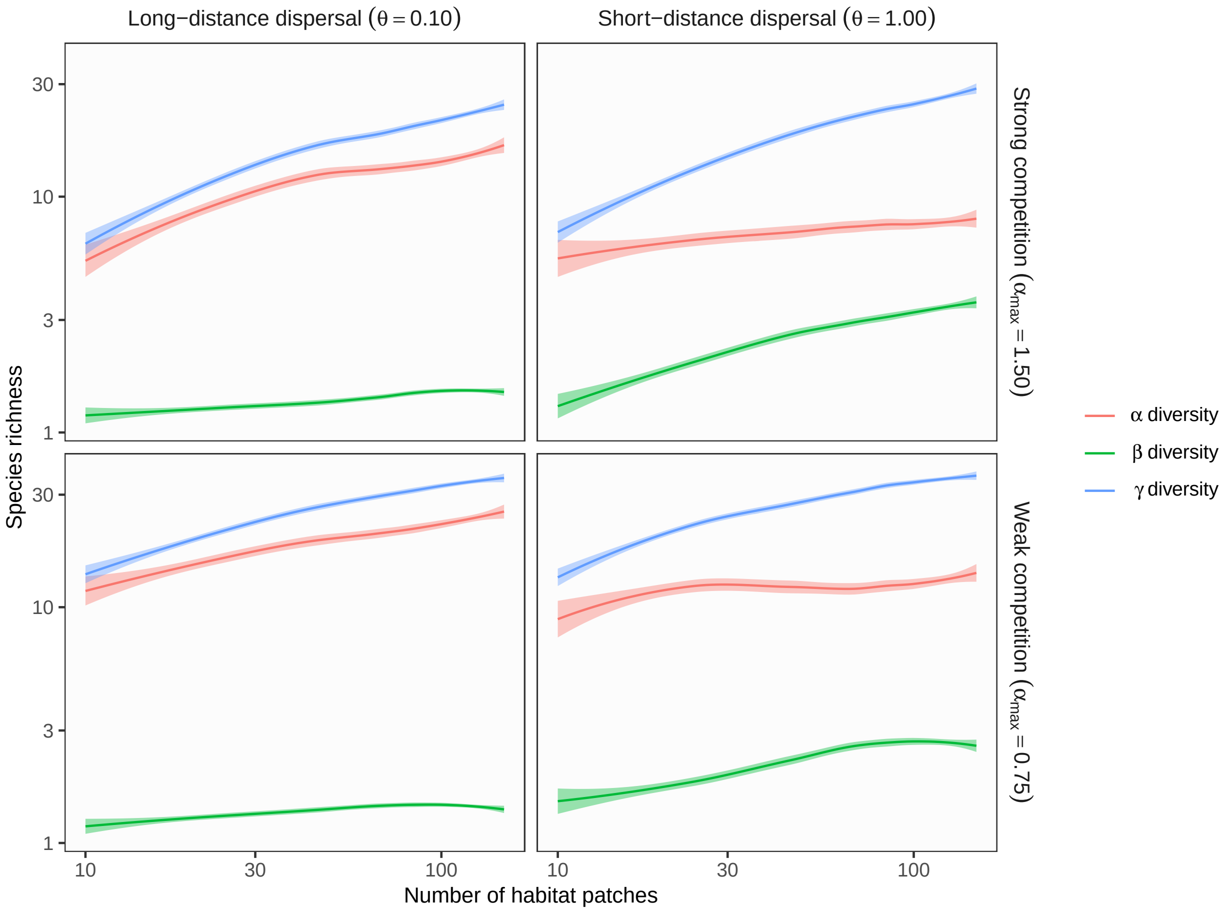

**Figure S6** Theoretical predictions for ecosystem size influences (the number of habitat patches) on $\alpha$, $\beta$, and $\gamma$ diversity in branching networks. In this simulation, environmental variation at headwaters ($\sigma_h$) is equal to local environmental noise ($\sigma_l$). Lines and shades are loess curves fitted to simulated data and their 95% confidence intervals. Each panel represents different ecological scenarios under which metacommunity dynamics were simulated. Rows represent different competition strength. Competitive coefficients ($\alpha_{ij}$) were varied randomly from 0 to 1.5 (top, strong competition) or 0.75 (bottom, weak competition). Columns represent different dispersal scenarios. Two dispersal parameters were chosen to simulate scenarios with long-distance (the rate parameter of an exponential dispersal kernel $\theta = 0.10$) and short-distance dispersal ($\theta = 1.0$). Other parameters are as follows: dispersal probability $p_d = 0.1$; environmental variation at headwaters $\sigma_h = 1$; local environmental noise $\sigma_l = 1$.

**Figure S7 Influence of ecosystem size** ($p_d = 0.1$, $\sigma_h = 0.01$, $\sigma_l = 0.01$)

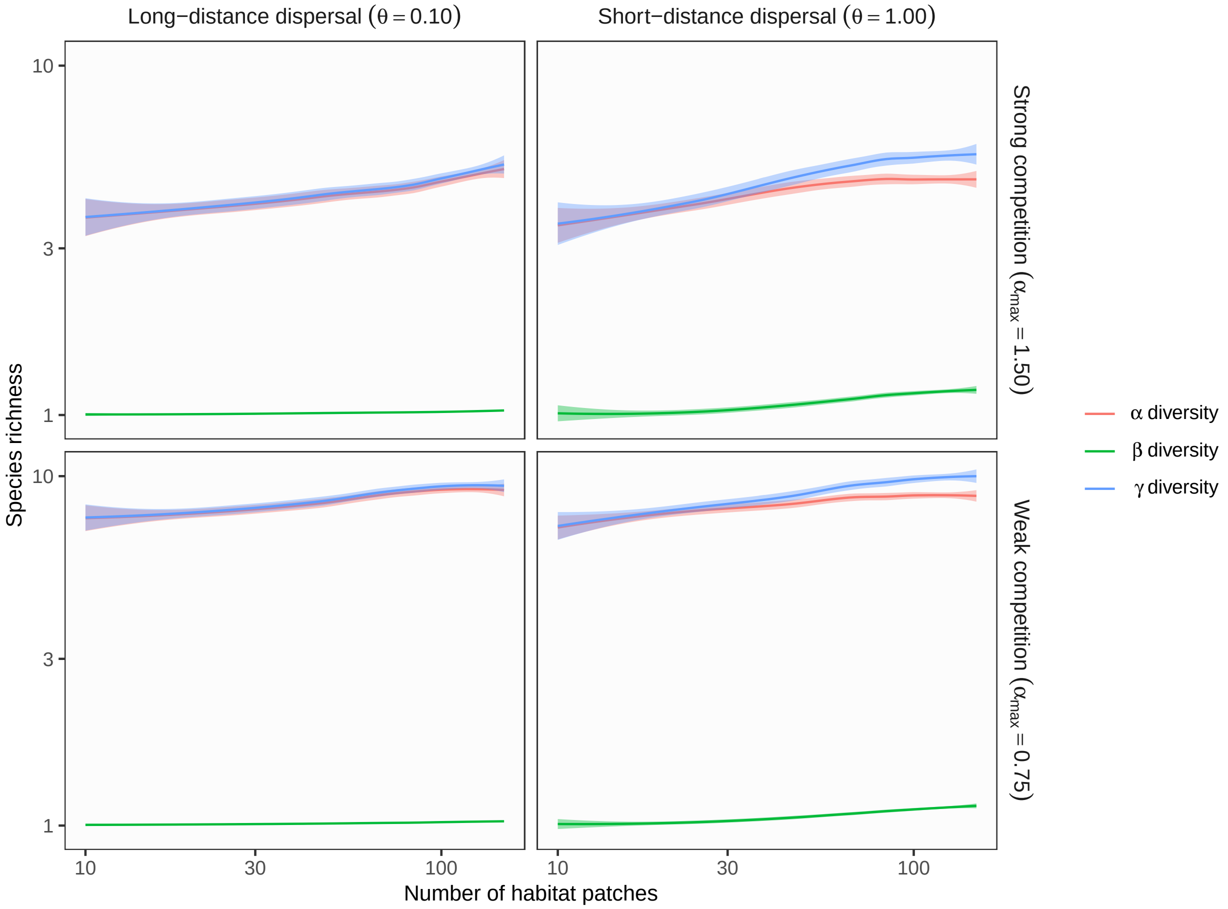

**Figure S7** Theoretical predictions for ecosystem size influences (the number of habitat patches) on $\alpha$, $\beta$, and $\gamma$ diversity in branching networks. In this simulation, environmental variation at headwaters ($\sigma_h$) is equal to local environmental noise ($\sigma_l$). Lines and shades are loess curves fitted to simulated data and their 95% confidence intervals. Each panel represents different ecological scenarios under which metacommunity dynamics were simulated. Rows represent different competition strength. Competitive coefficients ($\alpha_{ij}$) were varied randomly from 0 to 1.5 (top, strong competition) or 0.75 (bottom, weak competition). Columns represent different dispersal scenarios. Two dispersal parameters were chosen to simulate scenarios with long-distance (the rate parameter of an exponential dispersal kernel $\theta = 0.10$) and short-distance dispersal ($\theta = 1.0$). Other parameters are as follows: dispersal probability $p_d = 0.1$; environmental variation at headwaters $\sigma_h = 0.01$; local environmental noise $\sigma_l = 0.01$.

**Figure S8 Influence of ecosystem size** ($p_d = 0.1$, $\sigma_h = 0.01$, $\sigma_l = 1$)

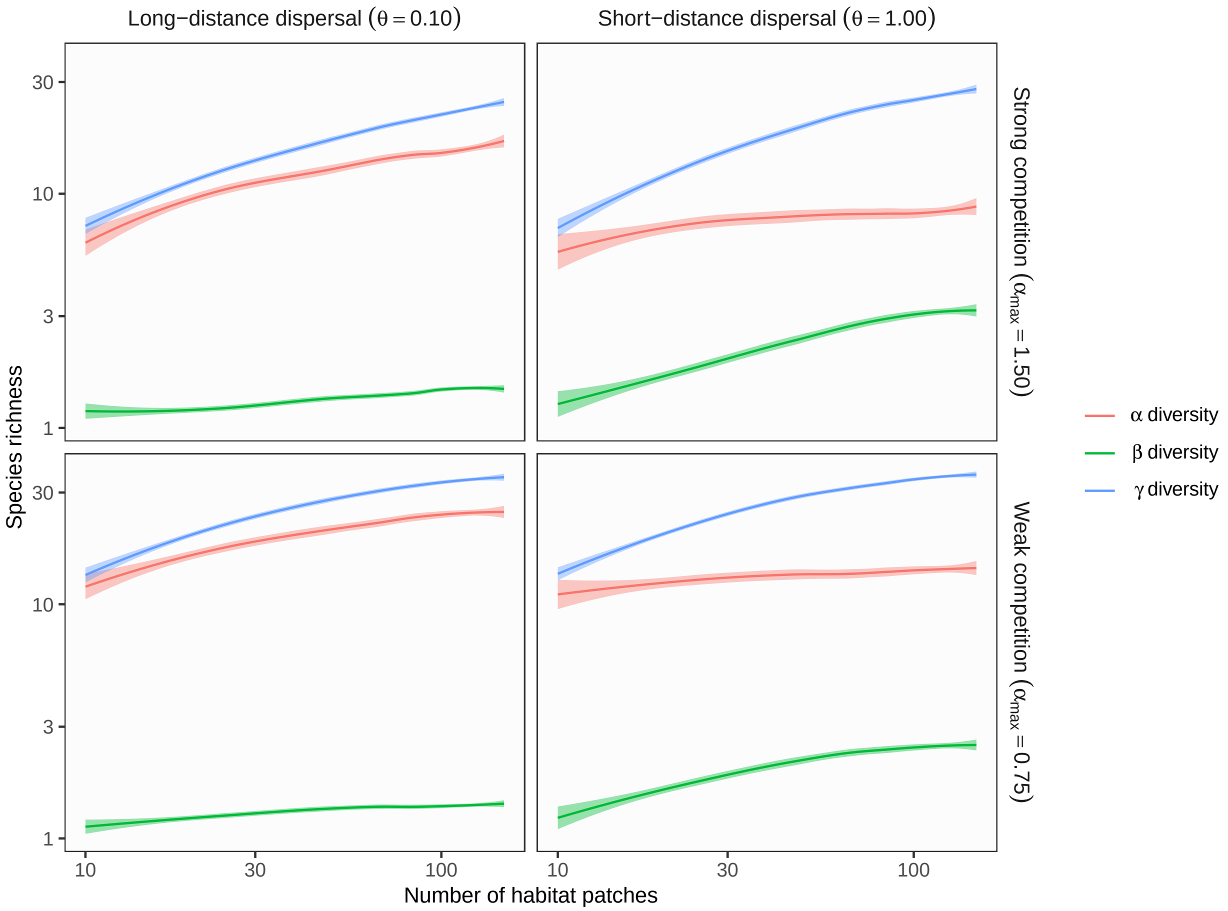

**Figure S8** Theoretical predictions for ecosystem size influences (the number of habitat patches) on $\alpha$, $\beta$, and $\gamma$ diversity in branching networks. In this simulation, environmental variation at headwaters ($\sigma_h$) is less than local environmental noise ($\sigma_l$). Lines and shades are loess curves fitted to simulated data and their 95% confidence intervals. Each panel represents different ecological scenarios under which metacommunity dynamics were simulated. Rows represent different competition strength. Competitive coefficients ($\alpha_{ij}$) were varied randomly from 0 to 1.5 (top, strong competition) or 0.75 (bottom, weak competition). Columns represent different dispersal scenarios. Two dispersal parameters were chosen to simulate scenarios with long-distance (the rate parameter of an exponential dispersal kernel $\theta = 0.10$) and short-distance dispersal ($\theta = 1.0$). Other parameters are as follows: dispersal probability $p_d = 0.1$; environmental variation at headwaters $\sigma_h = 0.01$; local environmental noise $\sigma_l = 1$.

**Figure S9 Influence of ecosystem size** ($p_d = 0.01$, $\sigma_h = 1$, $\sigma_l = 1$)

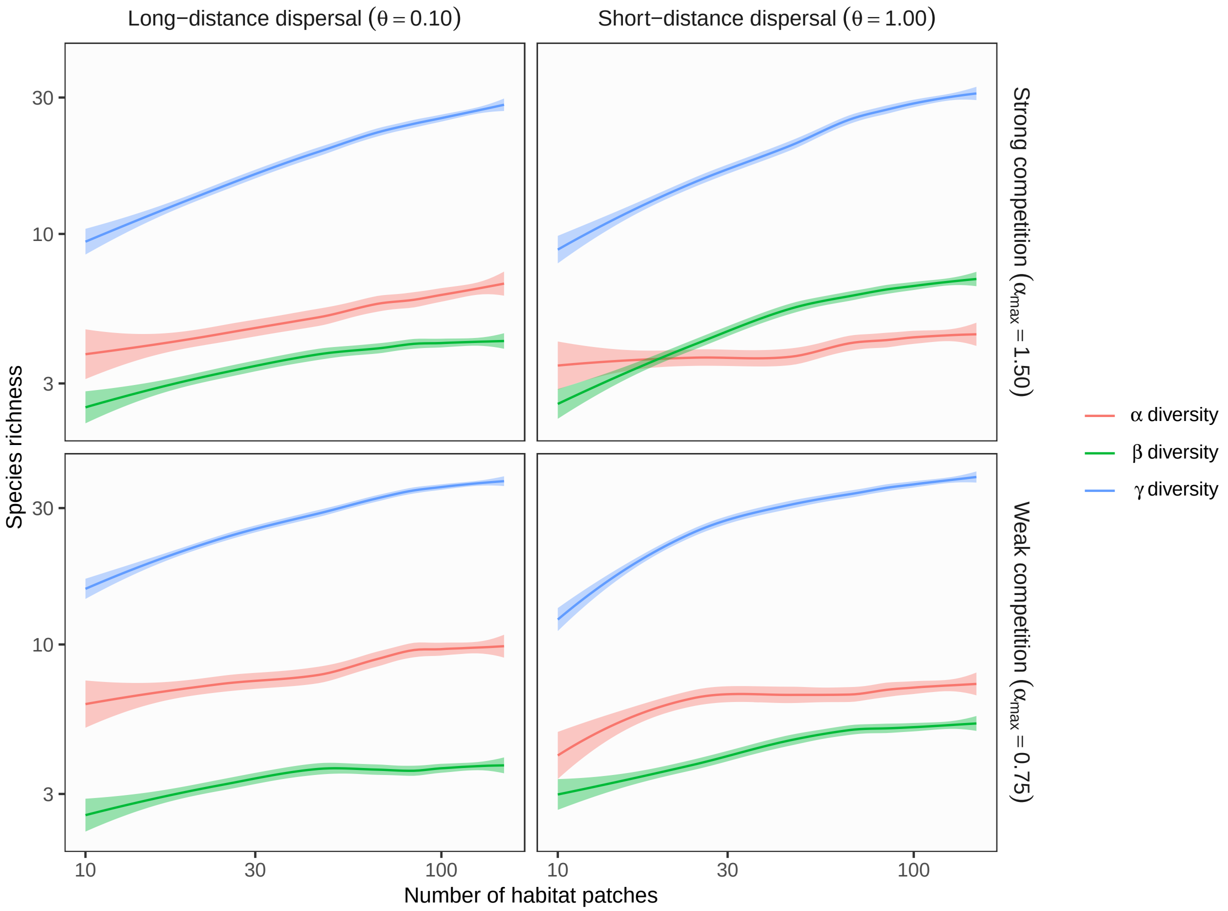

**Figure S9** Theoretical predictions for ecosystem size influences (the number of habitat patches) on $\alpha$, $\beta$, and $\gamma$ diversity in branching networks. In this simulation, environmental variation at headwaters ($\sigma_h$) is equal to local environmental noise ($\sigma_l$). Lines and shades are loess curves fitted to simulated data and their 95% confidence intervals. Each panel represents different ecological scenarios under which metacommunity dynamics were simulated. Rows represent different competition strength. Competitive coefficients ($\alpha_{ij}$) were varied randomly from 0 to 1.5 (top, strong competition) or 0.75 (bottom, weak competition). Columns represent different dispersal scenarios. Two dispersal parameters were chosen to simulate scenarios with long-distance (the rate parameter of an exponential dispersal kernel $\theta = 0.10$) and short-distance dispersal ($\theta = 1.0$). Other parameters are as follows: dispersal probability $p_d = 0.01$; environmental variation at headwaters $\sigma_h = 1$; local environmental noise $\sigma_l = 1$.

**Figure S10 Influence of ecosystem size** ($p_d = 0.01$, $\sigma_h = 0.01$, $\sigma_l = 0.01$)

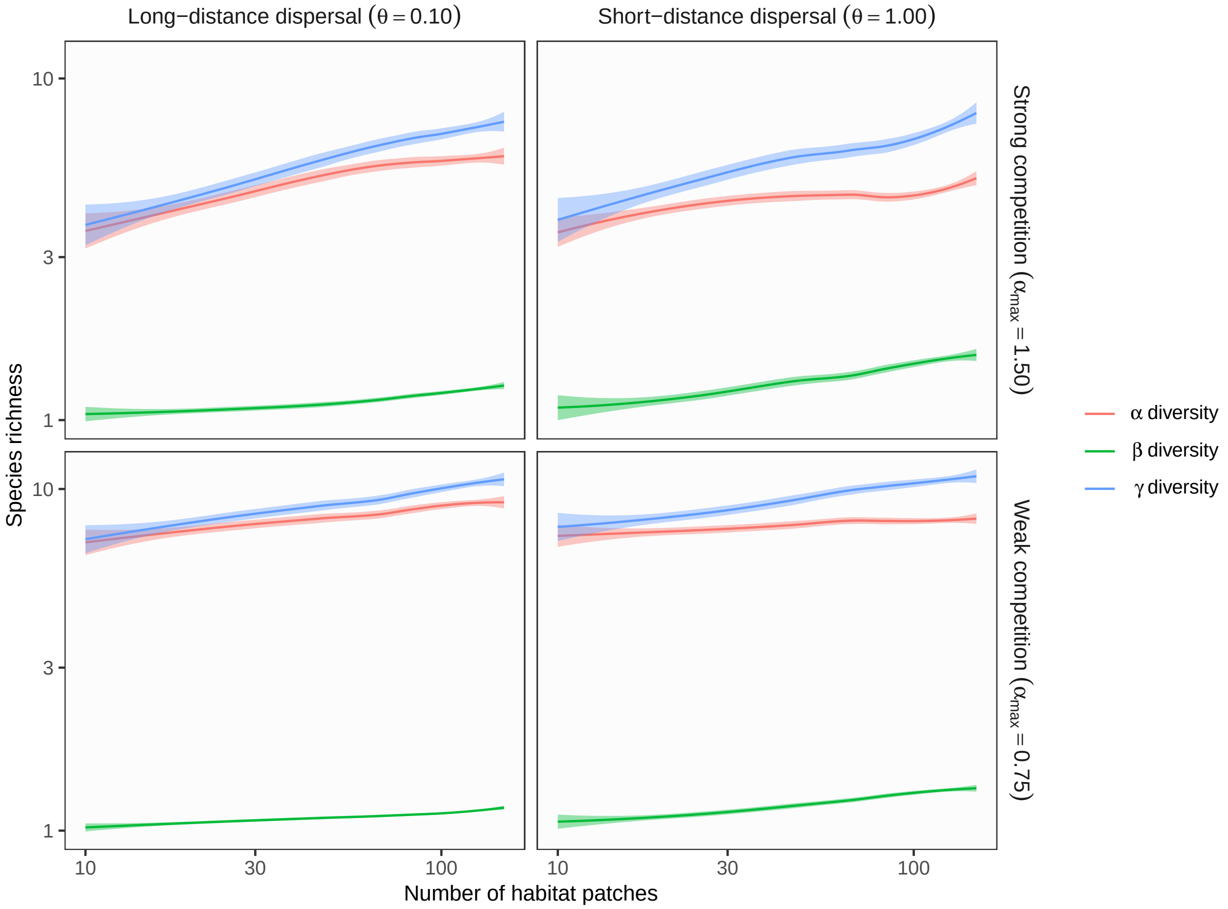

**Figure S10** Theoretical predictions for ecosystem size influences (the number of habitat patches) on $\alpha$, $\beta$, and $\gamma$ diversity in branching networks. In this simulation, environmental variation at headwaters ($\sigma_h$) is equal to local environmental noise ($\sigma_l$). Lines and shades are loess curves fitted to simulated data and their 95% confidence intervals. Each panel represents different ecological scenarios under which metacommunity dynamics were simulated. Rows represent different competition strength. Competitive coefficients ($\alpha_{ij}$) were varied randomly from 0 to 1.5 (top, strong competition) or 0.75 (bottom, weak competition). Columns represent different dispersal scenarios. Two dispersal parameters were chosen to simulate scenarios with long-distance (the rate parameter of an exponential dispersal kernel $\theta = 0.10$) and short-distance dispersal ($\theta = 1.0$). Other parameters are as follows: dispersal probability $p_d = 0.01$; environmental variation at headwaters $\sigma_h = 0.01$; local environmental noise $\sigma_l = 0.01$.

**Figure S11 Influence of ecosystem size** ($p_d = 0.01$, $\sigma_h = 0.01$, $\sigma_l = 1$)

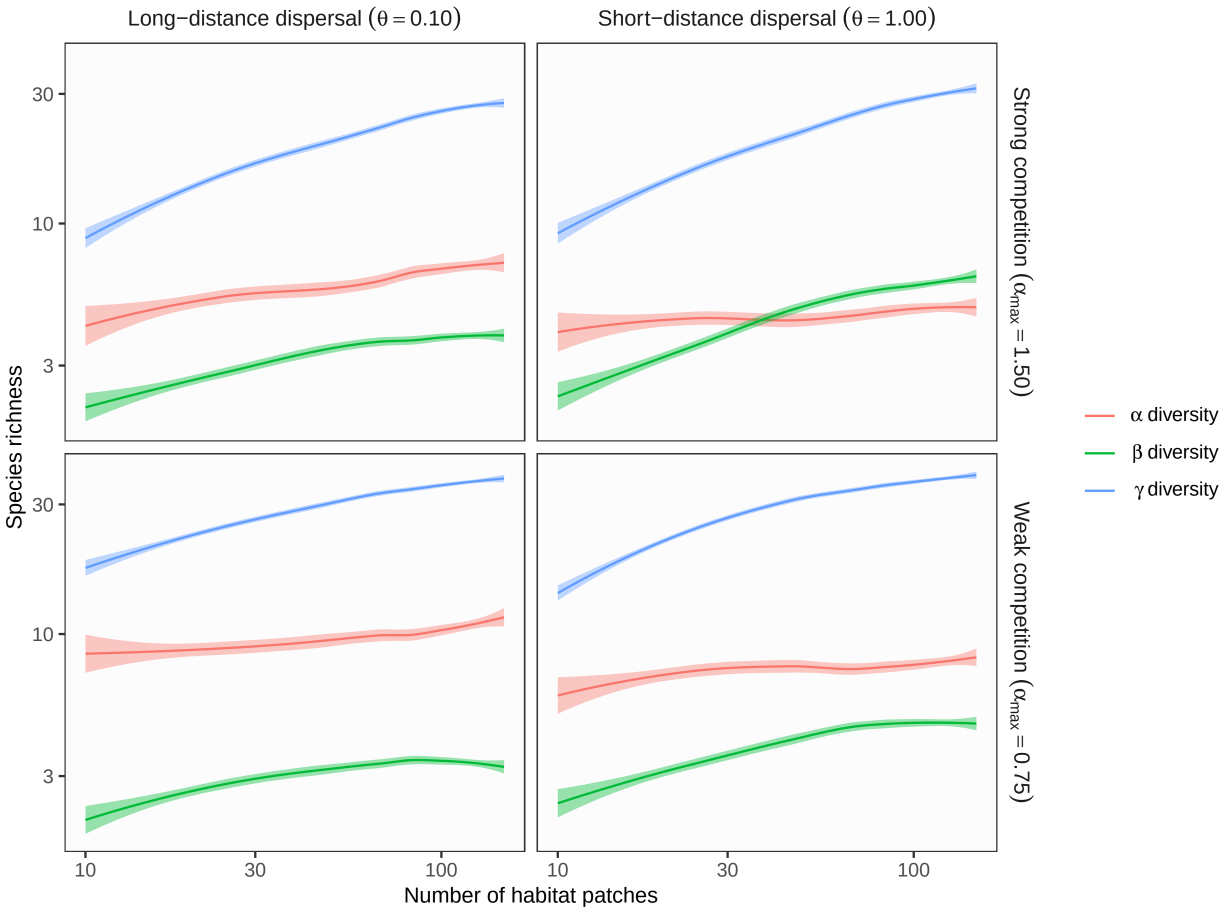

**Figure S11** Theoretical predictions for ecosystem size influences (the number of habitat patches) on $\alpha$, $\beta$, and $\gamma$ diversity in branching networks. In this simulation, environmental variation at headwaters ($\sigma_h$) is less than local environmental noise ($\sigma_l$). Lines and shades are loess curves fitted to simulated data and their 95% confidence intervals. Each panel represents different ecological scenarios under which metacommunity dynamics were simulated. Rows represent different competition strength. Competitive coefficients ($\alpha_{ij}$) were varied randomly from 0 to 1.5 (top, strong competition) or 0.75 (bottom, weak competition). Columns represent different dispersal scenarios. Two dispersal parameters were chosen to simulate scenarios with long-distance (the rate parameter of an exponential dispersal kernel $\theta = 0.10$) and short-distance dispersal ($\theta = 1.0$). Other parameters are as follows: dispersal probability $p_d = 0.01$; environmental variation at headwaters $\sigma_h = 0.01$; local environmental noise $\sigma_l = 1$.

**Figure S12 Influence of ecosystem complexity** ($p_d = 0.1$, $\sigma_h = 1$, $\sigma_l = 0.01$)

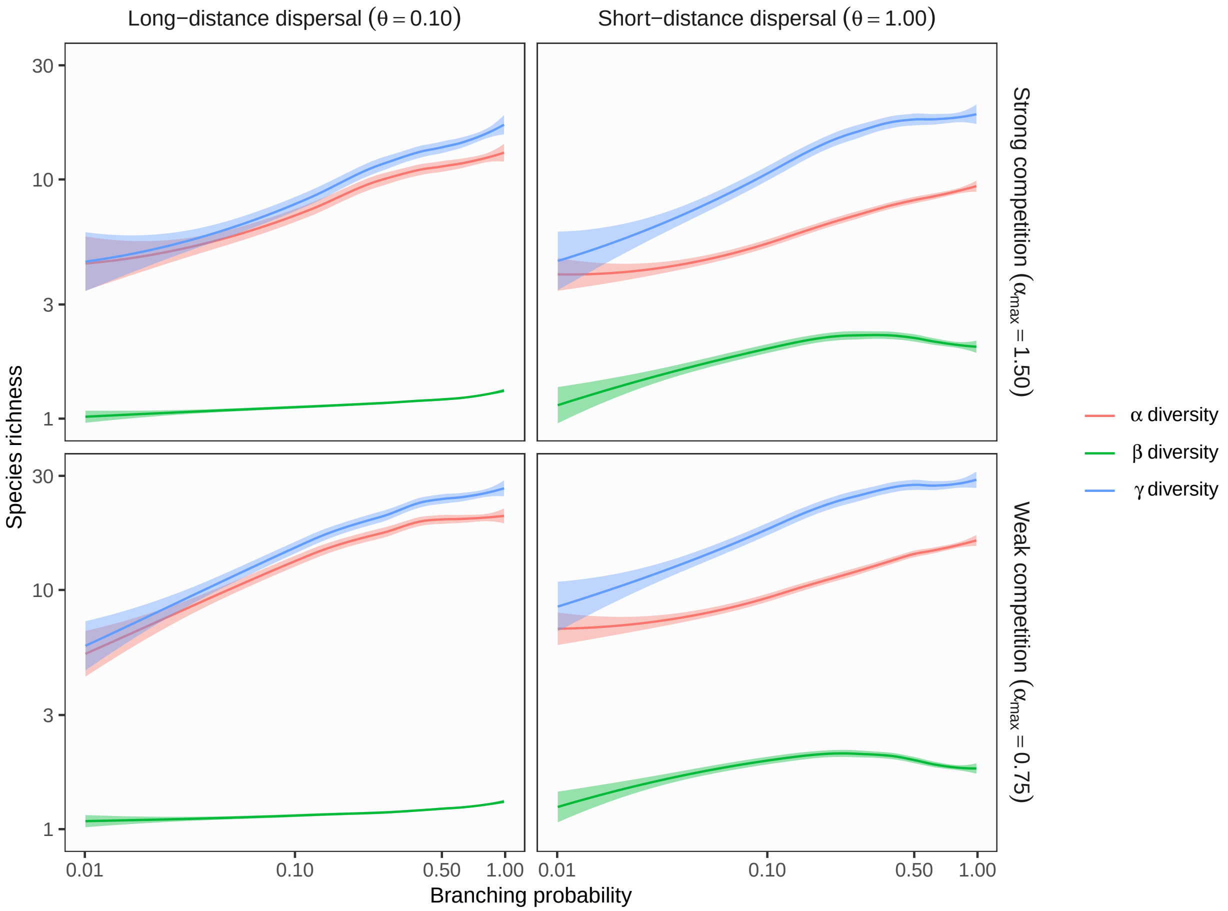

**Figure S12** Theoretical predictions for ecosystem complexity influences (branching probability) on $\alpha$, $\beta$, and $\gamma$ diversity in branching networks. In this simulation, environmental variation at headwaters ($\sigma_h$) exceeds local environmental noise ($\sigma_l$). Lines and shades are loess curves fitted to simulated data and their 95% confidence intervals. Each panel represents different ecological scenarios under which metacommunity dynamics were simulated. Rows represent different competition strength. Competitive coefficients ($\alpha_{ij}$) were varied randomly from 0 to 1.5 (top, strong competition) or 0.75 (bottom, weak competition). Columns represent different dispersal scenarios. Two dispersal parameters were chosen to simulate scenarios with long-distance (the rate parameter of an exponential dispersal kernel $\theta = 0.10$) and short-distance dispersal ($\theta = 1.0$). Other parameters are as follows: dispersal probability $p_d = 0.1$; environmental variation at headwaters $\sigma_h = 1$; local environmental noise $\sigma_l = 0.01$.

**Figure S13 Influence of ecosystem complexity** ($p_d = 0.1$, $\sigma_h = 1$, $\sigma_l = 1$)

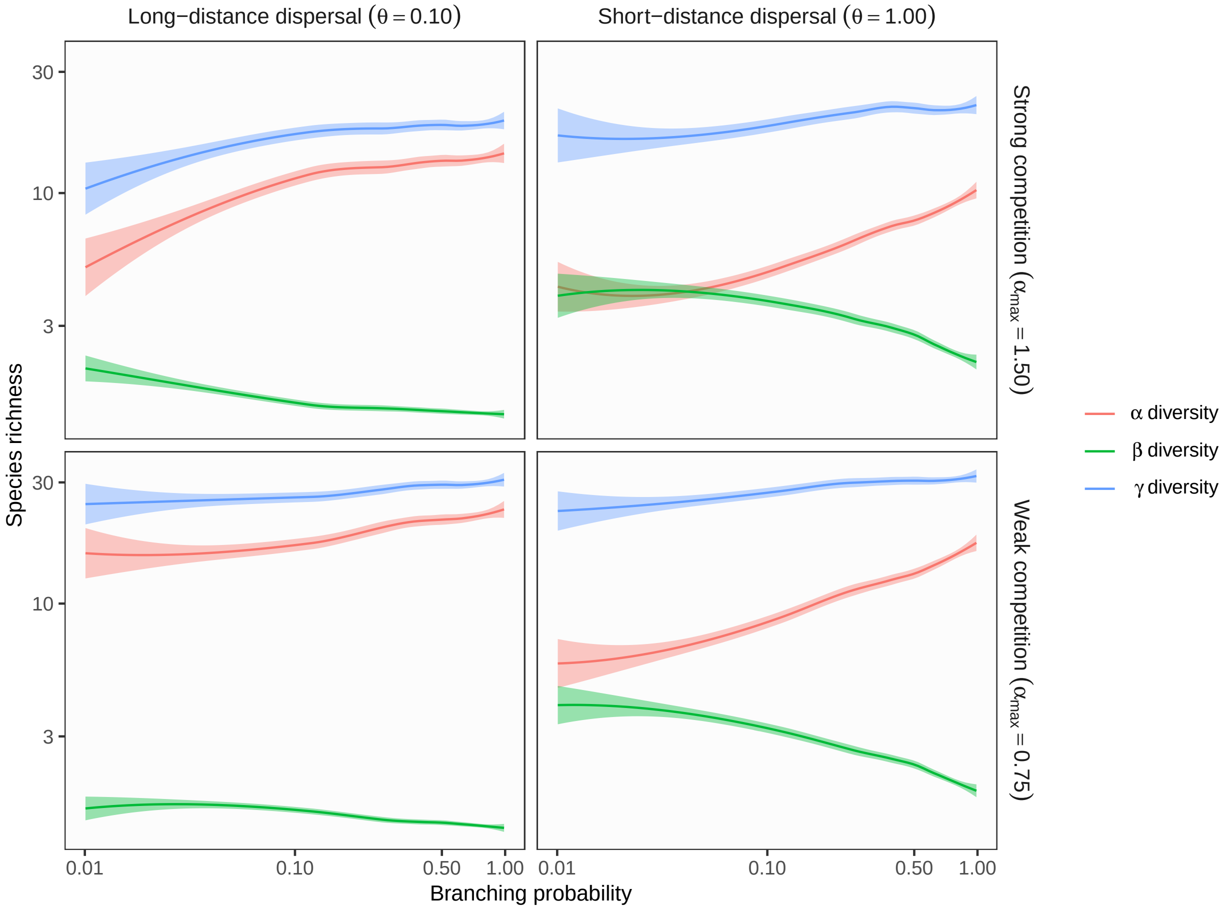

**Figure S13** Theoretical predictions for ecosystem complexity influences (branching probability) on $\alpha$, $\beta$, and $\gamma$ diversity in branching networks. In this simulation, environmental variation at headwaters ($\sigma_h$) is equal to local environmental noise ($\sigma_l$). Lines and shades are loess curves fitted to simulated data and their 95% confidence intervals. Each panel represents different ecological scenarios under which metacommunity dynamics were simulated. Rows represent different competition strength. Competitive coefficients ($\alpha_{ij}$) were varied randomly from 0 to 1.5 (top, strong competition) or 0.75 (bottom, weak competition). Columns represent different dispersal scenarios. Two dispersal parameters were chosen to simulate scenarios with long-distance (the rate parameter of an exponential dispersal kernel $\theta = 0.10$) and short-distance dispersal ($\theta = 1.0$). Other parameters are as follows: dispersal probability $p_d = 0.1$; environmental variation at headwaters $\sigma_h = 1$; local environmental noise $\sigma_l = 1$.

**Figure S14 Influence of ecosystem complexity** ($p_d = 0.1$, $\sigma_h = 0.01$, $\sigma_l = 0.01$)

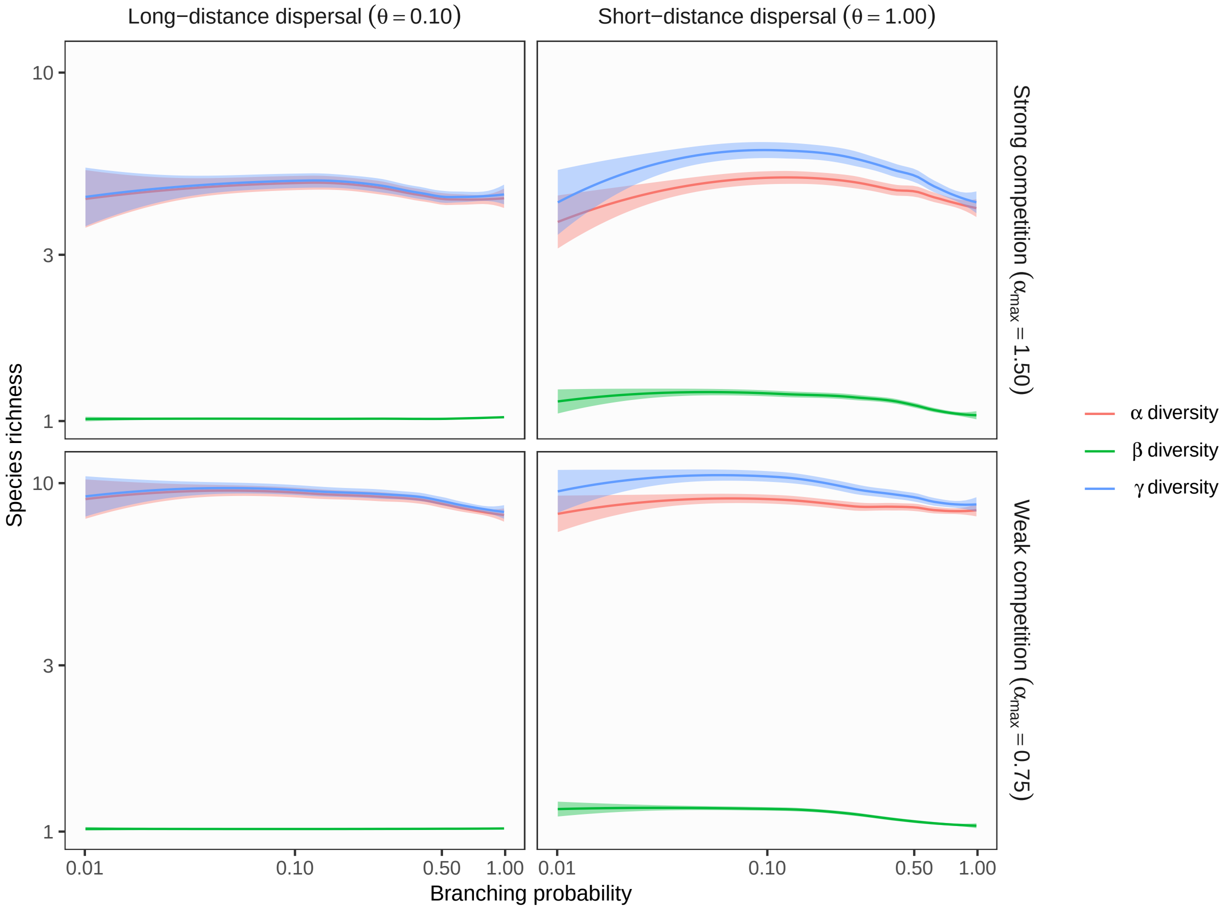

**Figure S14** Theoretical predictions for ecosystem complexity influences (branching probability) on $\alpha$, $\beta$, and $\gamma$ diversity in branching networks. In this simulation, environmental variation at headwaters ($\sigma_h$) is equal to local environmental noise ($\sigma_l$). Lines and shades are loess curves fitted to simulated data and their 95% confidence intervals. Each panel represents different ecological scenarios under which metacommunity dynamics were simulated. Rows represent different competition strength. Competitive coefficients ($\alpha_{ij}$) were varied randomly from 0 to 1.5 (top, strong competition) or 0.75 (bottom, weak competition). Columns represent different dispersal scenarios. Two dispersal parameters were chosen to simulate scenarios with long-distance (the rate parameter of an exponential dispersal kernel $\theta = 0.10$) and short-distance dispersal ($\theta = 1.0$). Other parameters are as follows: dispersal probability $p_d = 0.1$; environmental variation at headwaters $\sigma_h = 0.01$; local environmental noise $\sigma_l = 0.01$.

**Figure S15 Influence of ecosystem complexity** ($p_d = 0.1$, $\sigma_h = 0.01$, $\sigma_l = 1$)

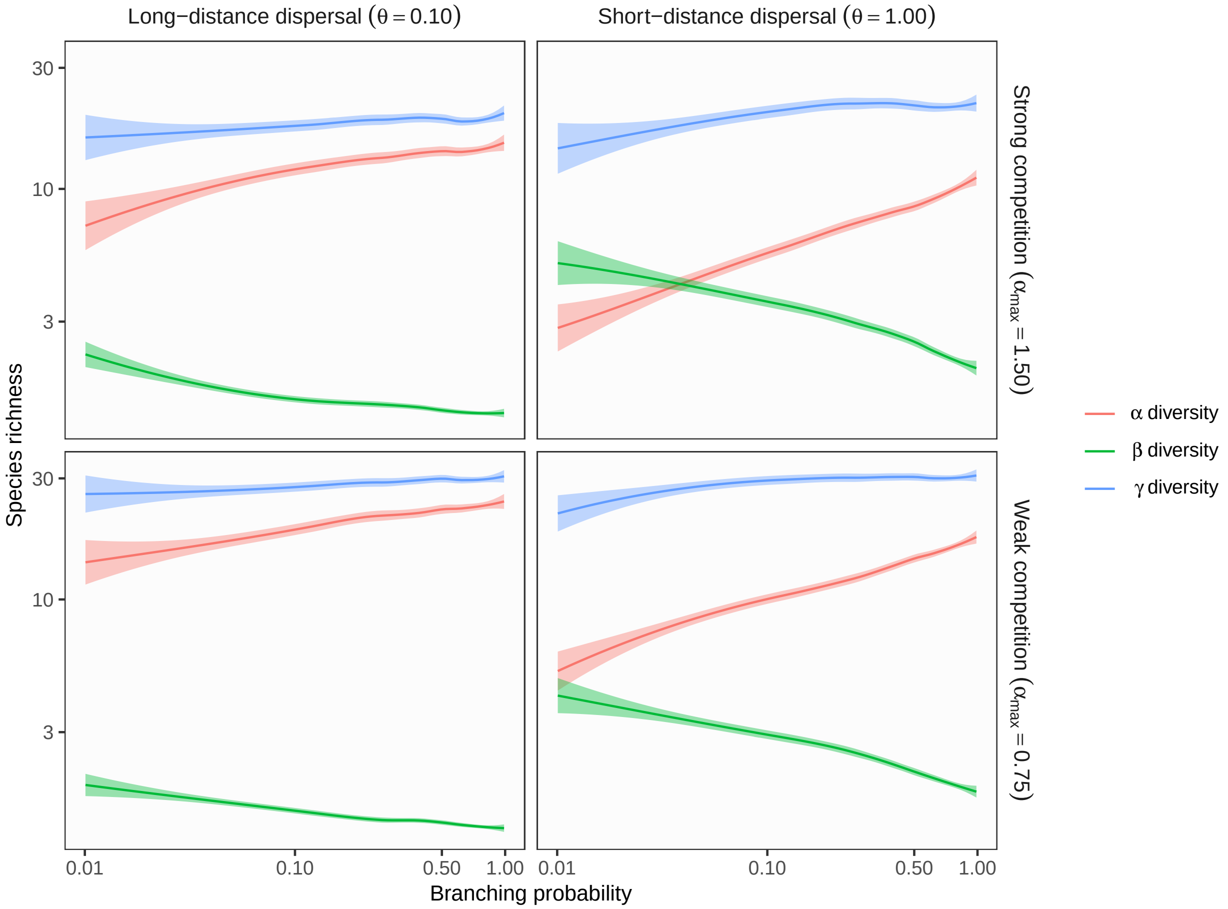

**Figure S15** Theoretical predictions for ecosystem complexity influences (branching probability) on $\alpha$, $\beta$, and $\gamma$ diversity in branching networks. In this simulation, environmental variation at headwaters ($\sigma_h$) is less than local environmental noise ($\sigma_l$). Lines and shades are loess curves fitted to simulated data and their 95% confidence intervals. Each panel represents different ecological scenarios under which metacommunity dynamics were simulated. Rows represent different competition strength. Competitive coefficients ($\alpha_{ij}$) were varied randomly from 0 to 1.5 (top, strong competition) or 0.75 (bottom, weak competition). Columns represent different dispersal scenarios. Two dispersal parameters were chosen to simulate scenarios with long-distance (the rate parameter of an exponential dispersal kernel $\theta = 0.10$) and short-distance dispersal ($\theta = 1.0$). Other parameters are as follows: dispersal probability $p_d = 0.1$; environmental variation at headwaters $\sigma_h = 0.01$; local environmental noise $\sigma_l = 1$.

**Figure S16 Influence of ecosystem complexity** ($p_d = 0.01$, $\sigma_h = 1$, $\sigma_l = 1$)

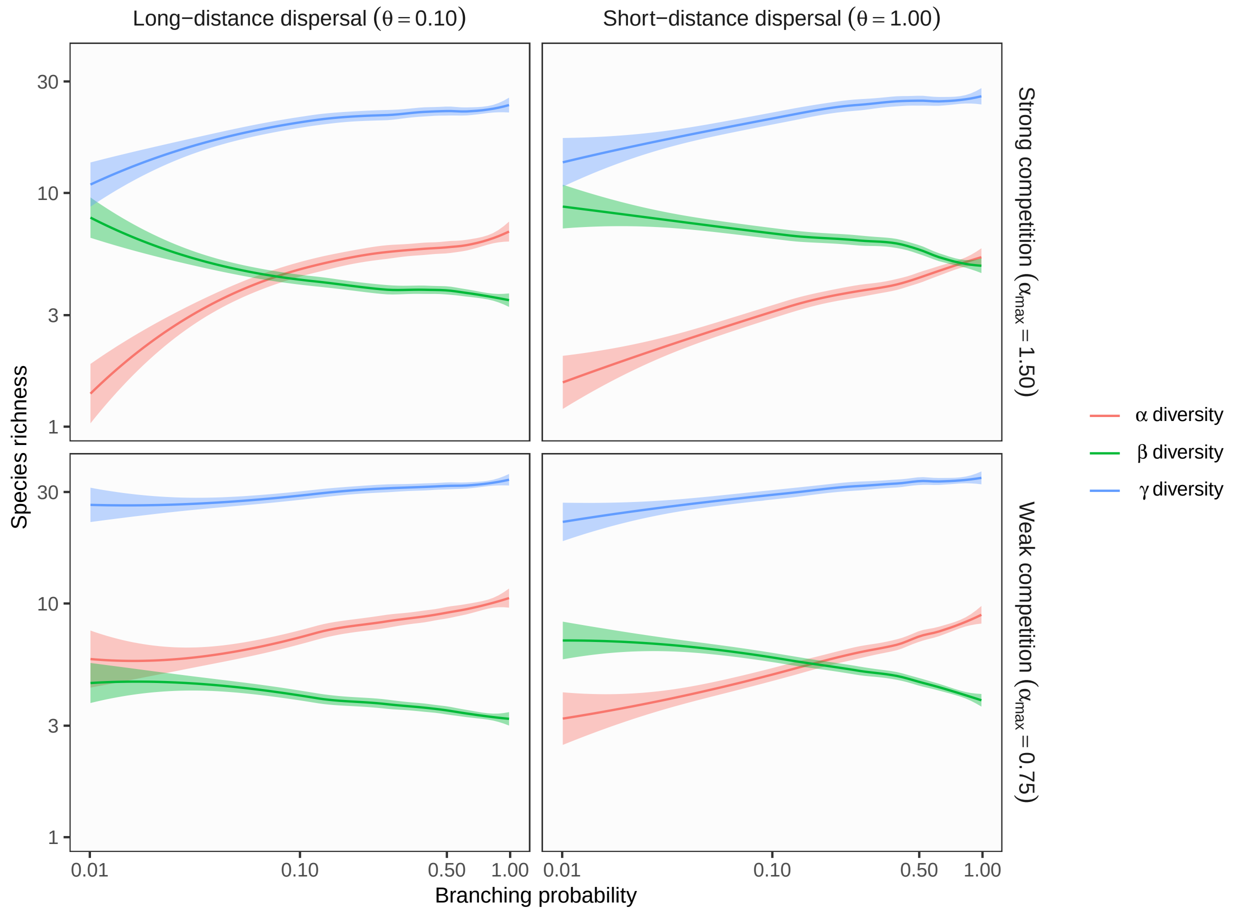

**Figure S16** Theoretical predictions for ecosystem complexity influences (branching probability) on $\alpha$, $\beta$, and $\gamma$ diversity in branching networks. In this simulation, environmental variation at headwaters ($\sigma_h$) is equal to local environmental noise ($\sigma_l$). Lines and shades are loess curves fitted to simulated data and their 95% confidence intervals. Each panel represents different ecological scenarios under which metacommunity dynamics were simulated. Rows represent different competition strength. Competitive coefficients ($\alpha_{ij}$) were varied randomly from 0 to 1.5 (top, strong competition) or 0.75 (bottom, weak competition). Columns represent different dispersal scenarios. Two dispersal parameters were chosen to simulate scenarios with long-distance (the rate parameter of an exponential dispersal kernel $\theta = 0.10$) and short-distance dispersal ($\theta = 1.0$). Other parameters are as follows: dispersal probability $p_d = 0.01$; environmental variation at headwaters $\sigma_h = 1$; local environmental noise $\sigma_l = 1$.

**Figure S17 Influence of ecosystem complexity** ($p_d = 0.01$, $\sigma_h = 0.01$, $\sigma_l = 0.01$)

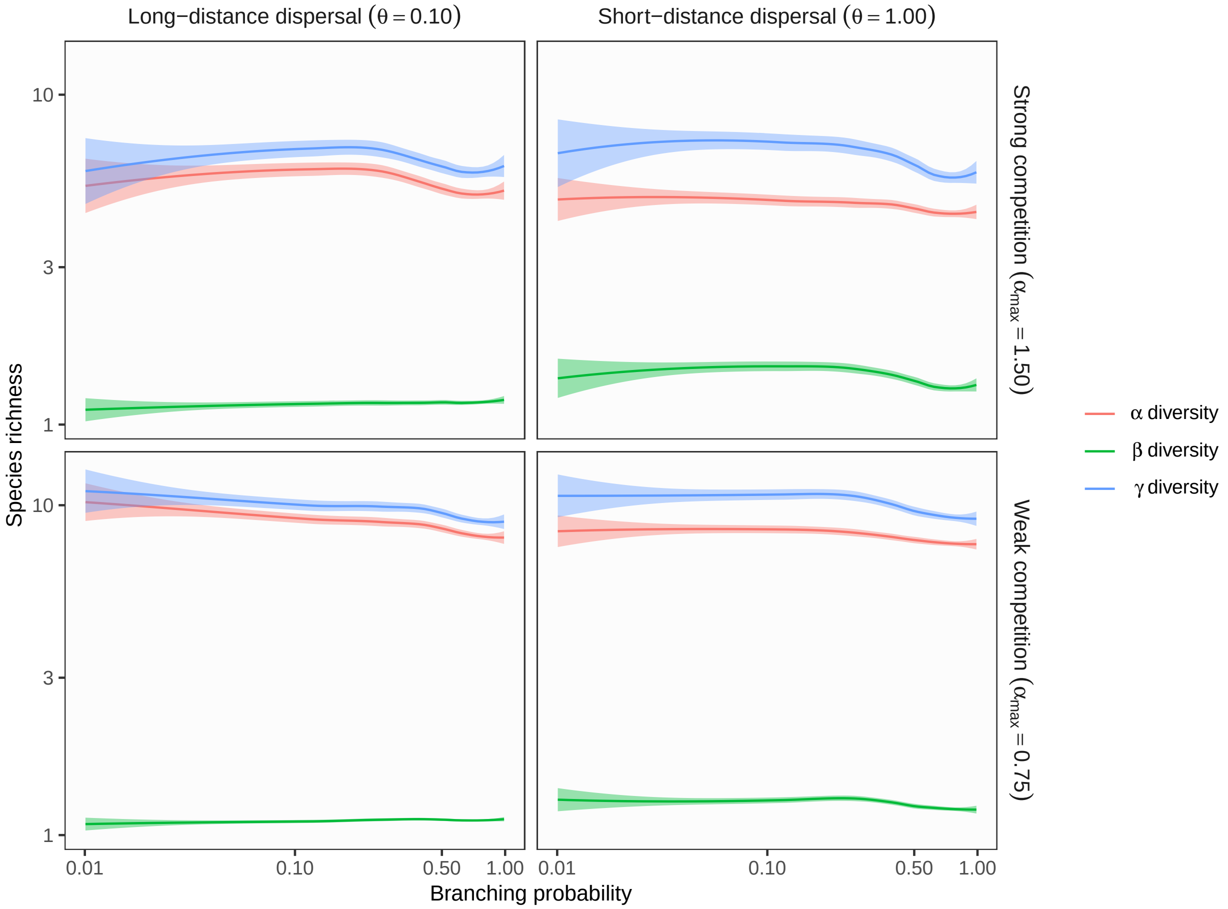

**Figure S17** Theoretical predictions for ecosystem complexity influences (branching probability) on $\alpha$, $\beta$, and $\gamma$ diversity in branching networks. In this simulation, environmental variation at headwaters ($\sigma_h$) is equal to local environmental noise ($\sigma_l$). Lines and shades are loess curves fitted to simulated data and their 95% confidence intervals. Each panel represents different ecological scenarios under which metacommunity dynamics were simulated. Rows represent different competition strength. Competitive coefficients ($\alpha_{ij}$) were varied randomly from 0 to 1.5 (top, strong competition) or 0.75 (bottom, weak competition). Columns represent different dispersal scenarios. Two dispersal parameters were chosen to simulate scenarios with long-distance (the rate parameter of an exponential dispersal kernel $\theta = 0.10$) and short-distance dispersal ($\theta = 1.0$). Other parameters are as follows: dispersal probability $p_d = 0.01$; environmental variation at headwaters $\sigma_h = 0.01$; local environmental noise $\sigma_l = 0.01$.

**Figure S18 Influence of ecosystem complexity** ($p_d = 0.01$, $\sigma_h = 0.01$, $\sigma_l = 1$)

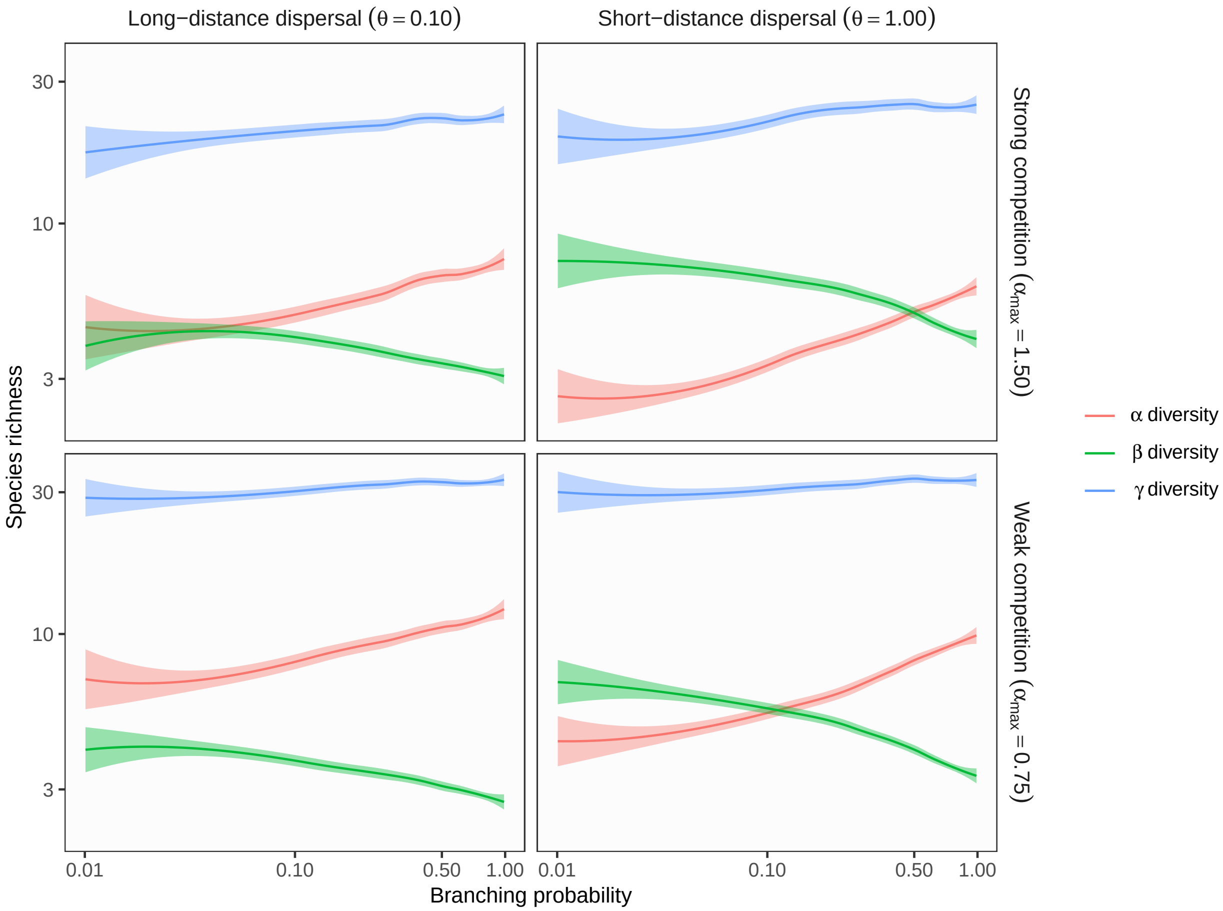

**Figure S18** Theoretical predictions for ecosystem complexity influences (branching probability) on $\alpha$, $\beta$, and $\gamma$ diversity in branching networks. In this simulation, environmental variation at headwaters ($\sigma_h$) is less than local environmental noise ($\sigma_l$). Lines and shades are loess curves fitted to simulated data and their 95% confidence intervals. Each panel represents different ecological scenarios under which metacommunity dynamics were simulated. Rows represent different competition strength. Competitive coefficients ($\alpha_{ij}$) were varied randomly from 0 to 1.5 (top, strong competition) or 0.75 (bottom, weak competition). Columns represent different dispersal scenarios. Two dispersal parameters were chosen to simulate scenarios with long-distance (the rate parameter of an exponential dispersal kernel $\theta = 0.10$) and short-distance dispersal ($\theta = 1.0$). Other parameters are as follows: dispersal probability $p_d = 0.01$; environmental variation at headwaters $\sigma_h = 0.01$; local environmental noise $\sigma_l = 1$.

**Figure S19 Correlation structure of explanatory variables**

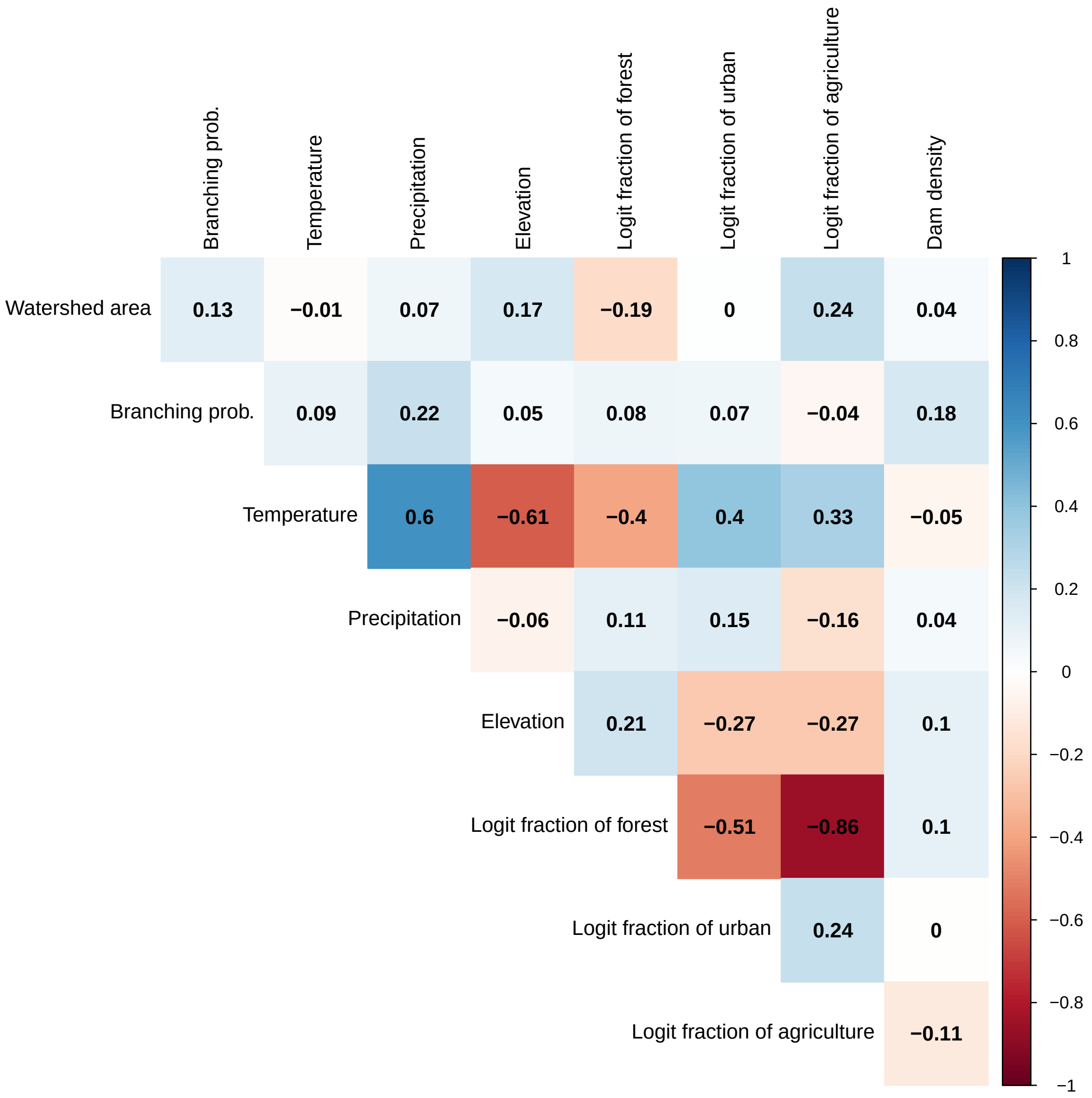

**Figure S19** Correlation structure of potential explanatory variables for riverine diversity metrics. Numeric values in each cell are the Pearson's correlation coefficients. Positive and negative correlations were colored in blue and red, respectively, and darker colors indicate stronger correlations. Environmental variables (temperature, precipitation, elevation, logit fraction of forest, logit fraction of urban, logit fraction of agriculture, and dam density) were expressed as deviations from the regional averages to remove any regional effects.

**Figure S20 Sensitivity analysis of asymptotic species richness**

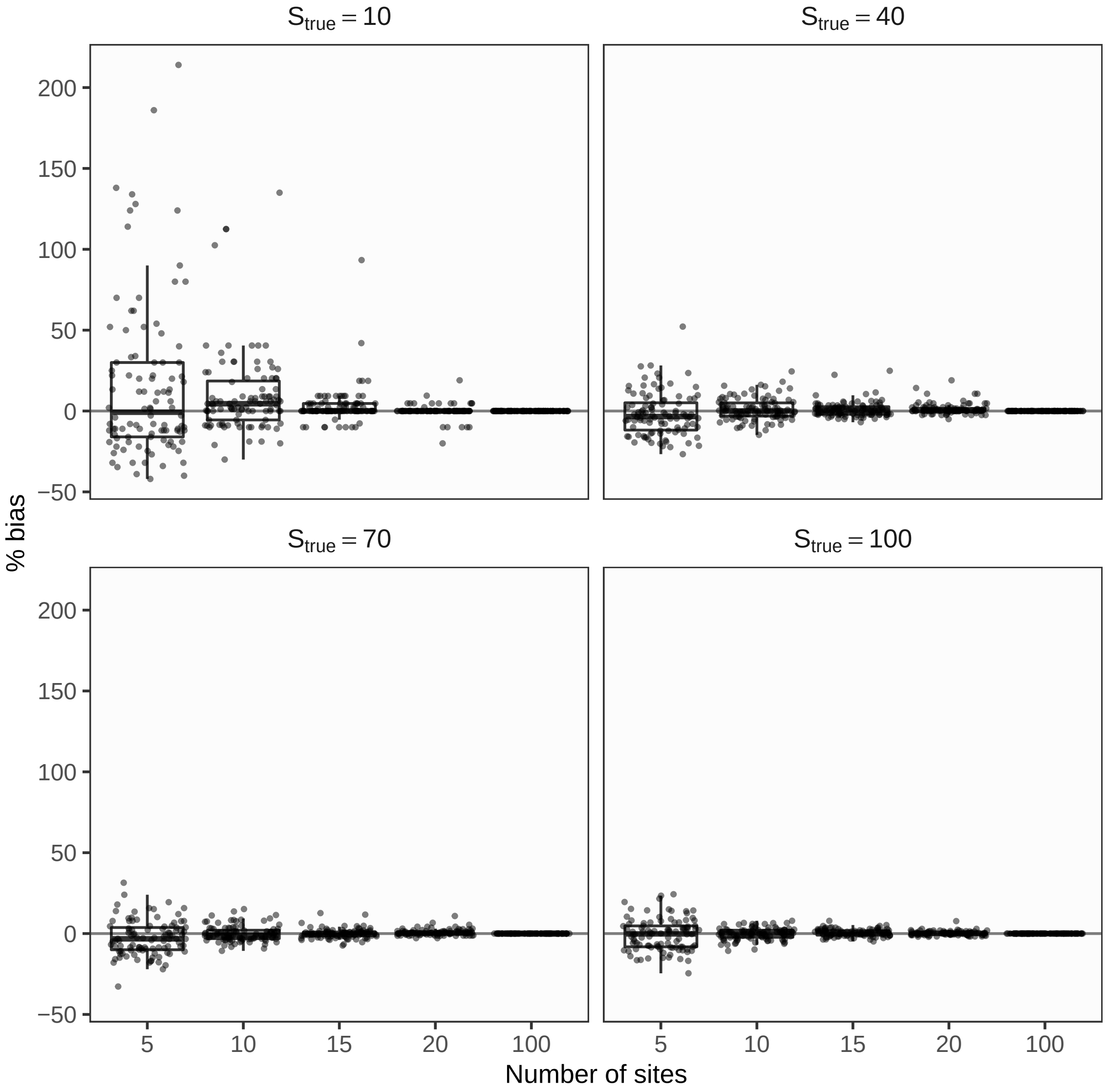

**Figure S20** Sensitivity analysis of asymptotic species richness in relation to true species richness $S_{true}$ (panels) and the number of sampling sites $N_{site}$ (x-axis). Positive values of % bias indicate an overestimation of species richness, while negative values indicate an underestimation. Different panels show results with different true species richness. The center lines are median values, the box boundaries 25 and 75 percentiles, and the whiskers 5 and 95 percentiles. Dots represent individual simulation replicates.
